# Expandable, Functional Hepatocytes Derived from Primary Cells Enable Liver Therapeutics

**DOI:** 10.1101/2024.12.28.630269

**Authors:** Sunil K. Mallanna, Soujanya S. Karanth, Joseph E. Marturano, Abhijith K. Kudva, Marcus Lehmann, Julie K. Morse, Morgan Jamiel, Timothy Norman, Christopher Wilson, Fabiola Munarin, David Broderick, Maxwell Van Buskirk, Esha Uddin, Michael Ret, Christopher Steele, Mehar Cheema, Justin Black, Eric Vanderploeg, Christopher Chen, Sangeeta Bhatia, Alireza Rezania, Thomas J. Lowery, Sophie Cazanave, Arnav Chhabra

**Affiliations:** Satellite Biosciences, Inc, Watertown, MA 02472, USA; Biological Design Center, Boston University, Boston, MA 02215, USA; Department of Biomedical Engineering, Boston University, Boston, MA 02215, USA; Wyss Institute for Biologically Inspired Engineering, Harvard University, Boston, MA 02215, USA; Institute for Medical Engineering and Science, Massachusetts Institute of Technology (MIT), Cambridge, MA 02139, USA; David H. Koch Institute for Integrative Cancer Research, MIT, Cambridge, MA 02139, USA; Broad Institute of MIT and Harvard, Cambridge, MA 02139, USA; Howard Hughes Medical Institute, Chevy Chase, MD 20815, USA

**Keywords:** liver cell therapy, hepatocyte expansion, liver disease, hepatocyte engraftment

## Abstract

Liver disease affects millions annually in the United States, with orthotopic transplantation as the only curative option for many patients. However, the scarcity of donor organs highlights a need for alternative cell-based therapies. Hepatocyte-based approaches are promising due to the cells’ inherent synthetic, metabolic, and detoxifying functions, but they face critical barriers, including the lack of a scalable source of functional hepatocytes and poor engraftment. In this study, we developed a scalable process for expanding primary human hepatocytes (PHHs) while preserving their identity and function. By leveraging heterocellular aggregation with stromal cells, we generated cryopreserved “seed” constructs that maintained viability and function post-thaw. Seeds demonstrated enhanced metabolic and detoxification functions and robust engraftment across multiple anatomic sites outside of the liver. Our approach addresses key limitations of hepatocyte-based therapies, offering a stable, scalable, and clinically viable platform for liver cell therapy applications.

## Introduction

The burden of liver disease in the United States is significant, affecting approximately 4.5 million patients and causing over 50,000 deaths annually (1). For many with liver failure or end-stage liver disease, liver transplantation remains the only curative option; however, a severe shortage of donor organs limits the accessibility of this life-saving treatment (2). This has spurred interest in developing alternative, cell-based therapies that can either supplement or replace the need for whole-organ transplants (2). Hepatocyte-based therapies are promising in this regard, as hepatocytes possess the synthetic, metabolic, and detoxifying functions required to restore liver function. However, progress has been hindered by two key challenges: (a) the lack of a sustainable source of functional hepatocytes, and (b) the poor engraftment observed in earlier clinical hepatocyte delivery attempts (3).

Over the past decade, various strategies have been explored to address these challenges. Primary human hepatocytes (PHHs) remain a potential cell type for this application due to their mature function, but their limited availability and high donor-to-donor variability hinder their widespread clinical use (3–6). Induced pluripotent stem cells (iPSCs) have emerged as a scalable option, as they can be differentiated into hepatocyte-like cells; however, these cells often exhibit immature functionality and lack the full metabolic capacity of PHHs (7, 8). Direct conversion of fibroblasts into hepatocyte-like cells has shown promise, but these cells engraft at low levels in vivo (9). Attempts to expand primary hepatocytes directly have led to some success but are often limited by loss of liver-specific functions or incompatibility with current Good Manufacturing Practices (cGMP) standards for clinical translation (10). In addition to sourcing functional cells, successful hepatocyte-based therapies must address the critical challenge of engraftment. Several groups have attempted to improve engraftment of hepatocytes by incorporating 3D culture conditions and supporting cells such as mesenchymal and endothelial cells (11–16). However, challenges remain, particularly with engraftment in fibrotic or extrahepatic environments (17), as well as scalability. These limitations highlight the need for innovative approaches that enable the scalable production of functional hepatocytes suitable for therapeutic use and strategies that allow hepatocytes to engraft efficiently at ectopic sites.

In this work, we aimed to address the limitations of hepatocyte-based therapies by evaluating whether primary hepatocytes can be expanded without dedifferentiation and whether heterocellular aggregation with supportive cells can improve engraftment at ectopic sites. Leveraging insights from developmental biology and liver regeneration, we developed a protocol to activate key regenerative pathways in PHHs, achieving robust expansion across multiple primary donors while preserving hepatic identity and function. We hypothesized that heterocellular aggregation with stromal support cells would create a more effective microenvironment for stability and engraftment (13, 18–20). To test this, we combined expanded hepatocytes with dermal fibroblasts in suspension bioreactors to generate “seed” constructs. We optimized cryopreservation protocols for seeds and assessed if the cryopreserved constructs retained comparable viability and function to freshly prepared ones. Finally, we evaluated engraftment at ectopic sites in rodents, including the spleen, fat pad, and kidney capsule, as proxies for clinically relevant extrahepatic locations. Our overarching goal was to establish an end-to-end, cGMP-compliant, scalable process for expanding PHHs and aggregating them with supportive cells to produce a stable, cryopreserved therapeutic product that can be shipped to the patient’s bedside for administration.

## Results

### Primary human hepatocytes from multiple donors can be expanded and scaled in culture

We have established an end-to-end workflow that begins with procuring donor livers and progresses through isolating and cryopreserving PHHs, expanding PHHs, forming heterocellular aggregates called “seeds,” cryopreserving the seeds, and finally transplanting them at ectopic sites (Fig. 1). We successfully isolated PHHs from cadaveric livers with high yield (>2E9) and post-thaw viability (>70%), exemplified by lots SAT-32 and SAT-33 (Fig. S1). Using insights from liver development and post-injury regeneration, we developed a primary hepatocyte expansion protocol incorporating factors that activate key regenerative pathways (Fig. 2A). This protocol enabled successful expansion from a small initial population of plated PHHs (Fig. S2) and proved effective across multiple donors, with 46% achieving greater than 20-fold expansion at passage 0 (P0) (Table 1). Interestingly, P0 expansion efficiency correlated negatively with donor age, explaining 42% of the variance in expansion potential (Fig. 2B).

**Figure 1:**
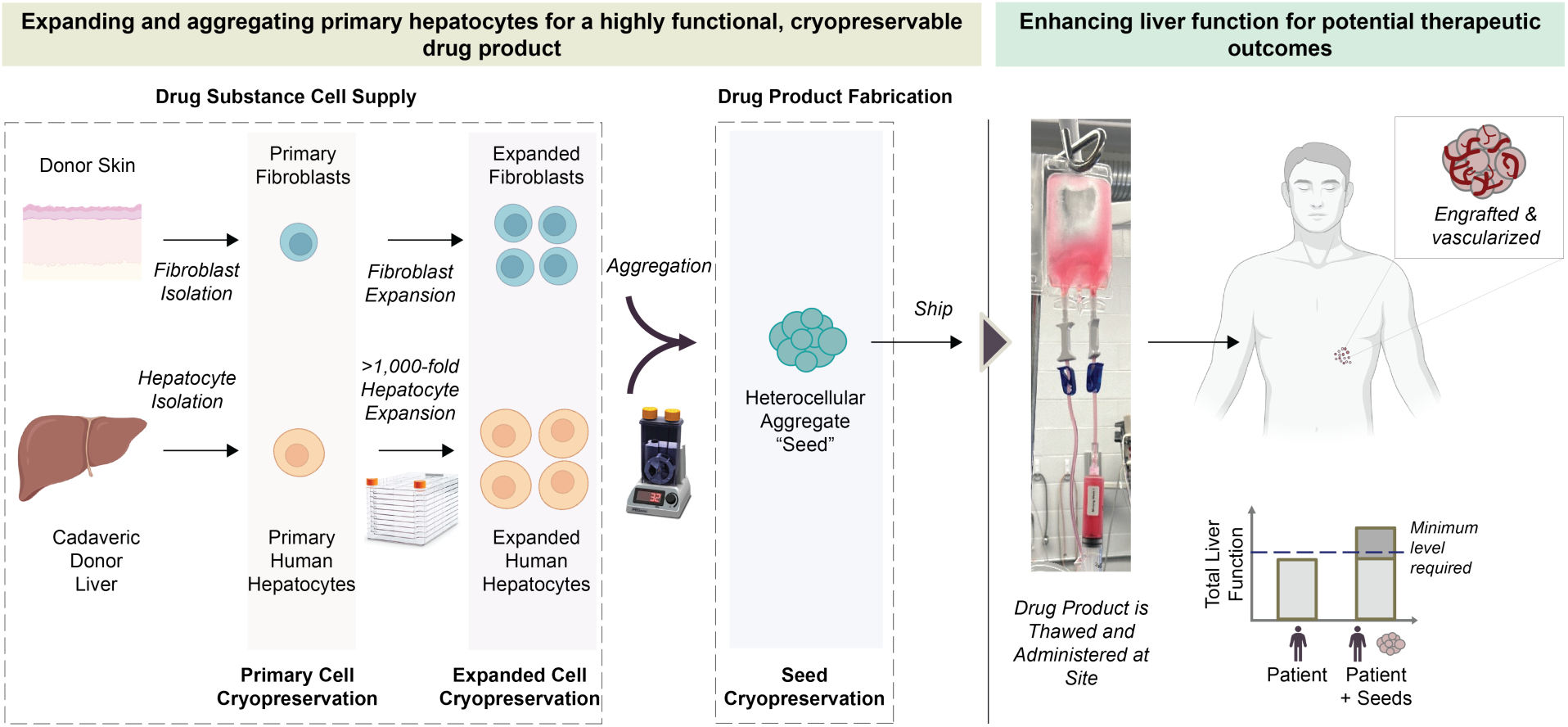
Schematic showing a cGMP-compatible process for creating a functional, cryopreserved drug product from primary hepatocytes, highlighting its potential therapeutic applications. Primary hepatocytes are isolated from donor livers, expanded in culture using CellStacks, and combined with dermal fibroblasts in bioreactors to form heterocellular aggregates, referred to as seeds. Seeds are stabilized through the support of extracellular matrix (ECM), paracrine signaling, and cell-to-cell interactions provided by the fibroblasts. The process is designed to decouple the generation of the drug substance and product, as both the cells and seeds are amenable to cryopreservation, and it is scalable for clinical product generation. The seeds can be administered directly or encapsulated in hydrogels to enhance liver function, providing a therapeutic approach for patients requiring liver support.

**Figure 2:**
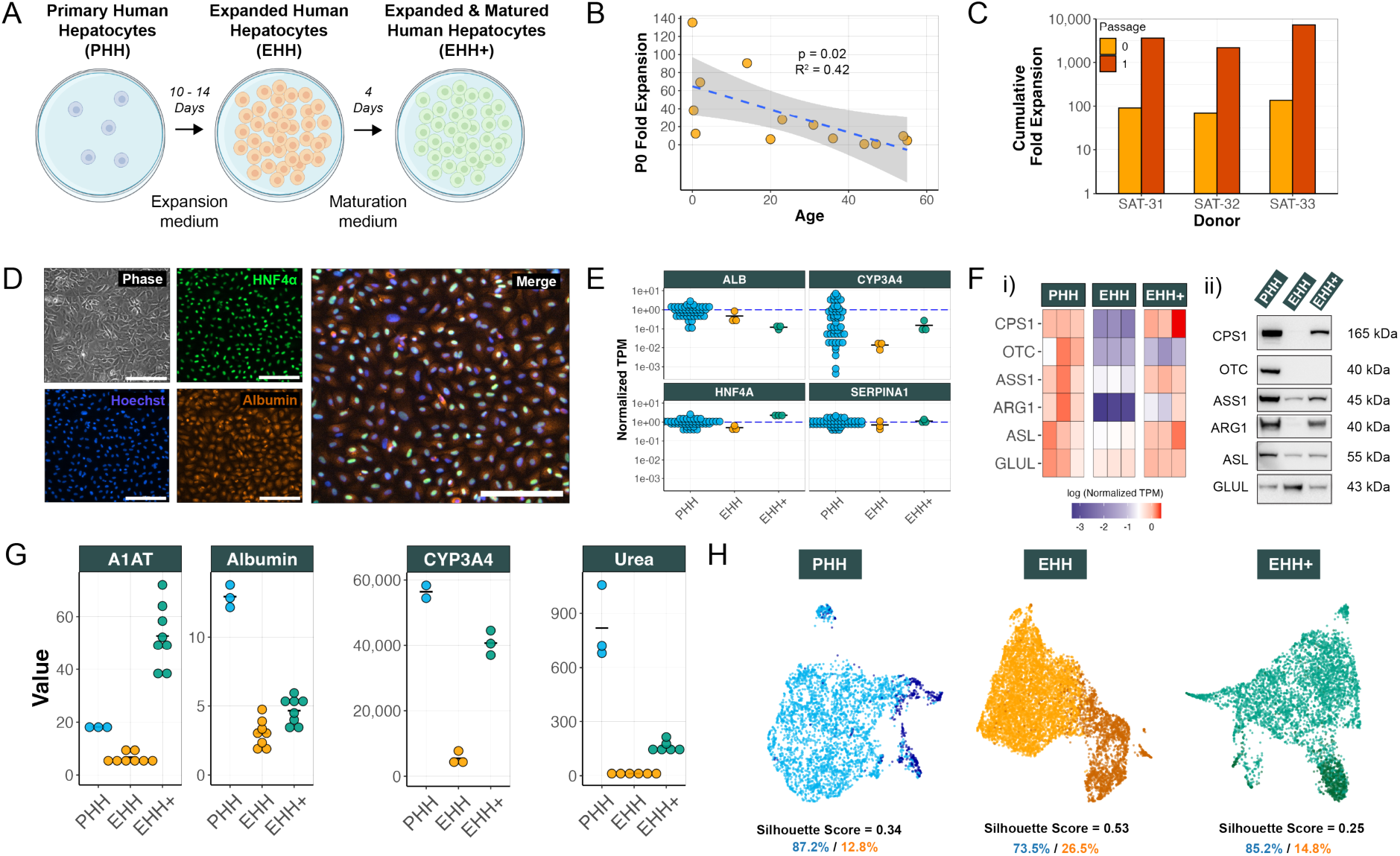
Primary hepatocyte expansion and maturation yield functional cell populations for therapeutic applications. (A) Illustration of hepatocyte culture, expansion, and maturation phases. (B) Correlation analysis showing a negative correlation between donor age and passage 0 (P0) fold expansion (p = 0.02, R^2^ = 0.42). (C) Cell count analysis demonstrating >2,000x cumulative fold expansion with a two-passage scheme across three donor lots. (D) Imaging analysis demonstrating the characteristic cuboidal morphology of expanded hepatocytes (Phase) and robust co-expression of HNF4α (green), albumin (orange), and Hoechst (nuclei). Over 99% of the expanded hepatocytes exhibited triple positivity, confirming their hepatic identity (scale bars = 200 μm). (E) Gene expression analysis showing comparable levels of liver-specific genes (ALB, CYP3A4, HNF4A, SERPINA1) across PHH, EHH, and EHH+ samples. Each dot represents a unique donor (n = 46 for PHH, n = 3 for EHH, n = 3 for EHH+). Transcripts per million (TPM) values are normalized to the mean of the PHH group. (F) (i) Gene expression and (ii) Western blot analysis of metabolic enzymes (CPS1, OTC, ASS1, ARG1, ASL, GLUL), showing decreased functional expression at the EHH stage with a subsequent restoration at the EHH+ stage. Each column in the heatmap represents a unique donor and TPM values are normalized to the mean TPM of the corresponding gene in the PHH group. (G) Functional assessment across synthetic (α1-antitrypsin, albumin) and metabolic (CYP3A4 induction, urea cycle) axes, demonstrating functional reduction at EHH stage with a partial restoration at the EHH+ stage. Units for A1AT, albumin and urea are µg/1E6 cells/day, while the units for CYP3A4 activity are relative luminescence units. (H) UMAP plots illustrating the transition from a homogeneous mature PHH population to two distinct sub-populations at the EHH stage: an expanding, immature group and a less expanding, mature group. This shifts back to a homogeneous mature population at the EHH+ stage. The percentages shown represent the proportion of each cell population.

**Table 1:** Fold expansion at passage 0 across multiple donors.

| **Donor** | **Fold expansion at passage 0** |
| --- | --- |
| SAT-21 | 6.13 |
| SAT-22 | 0.85 |
| SAT-23 | 4.65 |
| SAT-24 | 6.96 |
| SAT-25 | 0.78 |
| SAT-26 | 9.50 |
| SAT-27 | 21.90 |
| SAT-28 | 12.20 |
| SAT-29 | 27.75 |
| SAT-30 | 38.30 |
| SAT-31 | 90.50 |
| SAT-32 | 69.21 |
| SAT-33 | 135.33 |

**Table 2:** Demographic information for all primary human hepatocyte (PHH) donors. PHHs used in this study were obtained thanks to LifeNet Health, IIAM, Lonza Bioscience, BioIVT, Thermo Fisher Scientific, Yecuris, and Novabiosis.

| Age | Sex | Race | Cause of Death |
| --- | --- | --- | --- |
| 20 yo | Male | Caucasian | Cerebrovascular stroke |
| 47 yo | Male | Caucasian | Anoxia |
| 55 yo | Female | Caucasian | Cerebrovascular stroke |
| 36 yo | Male | African American | Head trauma |
| 44 yo | Female | Caucasian | Anoxia |
| 54 yo | Male | Caucasian | Anoxia |
| 31 yo | Female | Caucasian | Cerebrovascular stroke |
| 10 mo | Female | Black/Caucasian | Anoxia |
| 23 yo | Female | Caucasian | Cardiac arrest |
| 4 mo | Female | Caucasian | SIDS |
| 14 yo | Female | Hispanic | Head trauma |
| 2 yo | Female | Caucasian | Cerebrovascular stroke |
| 5 days | Female | Caucasian | Natural causes |
| 45 yo | Male | Caucasian | Anoxia 2nd to cardiovascular |
| 42 yo | Female | Caucasian | Anoxia 2nd to drug intoxication |
| 33 yo | Male | Caucasian | Drug intoxication |
| 50 yo | Male | Caucasian | Cerebrovascular stroke |
| 31 yo | Female | Asian | Anoxia |
| 38 yo | Male | Caucasian | Gunshot wound |
| 46 yo | Female | African American | Anoxia |
| 7 yo | Male | Caucasian | Anoxia |
| 63 yo | Female | African American | Cerebrovascular stroke |
| 3 yo | Male | Caucasian | Anoxia |
| 6 mo | Male | Hispanic | Anoxia |
| 37 yo | Male | Caucasian | Cerebrovascular stroke |
| 60 yo | Male | Caucasian | Anoxia |
| 37 yo | Female | Caucasian | Anoxia |
| 32 yo | Male | Hispanic | Cerebrovascular stroke |
| 40 yo | Male | Caucasian | Unknown |
| 20 yo | Male | Caucasian | Unknown |
| 20 yo | Female | Caucasian | Unknown |
| 33 yo | Female | Caucasian | Unknown |
| 49 yo | Male | Caucasian | Unknown |
| 56 yo | Female | Caucasian | Unknown |
| 54 yo | Male | Caucasian | Unknown |
| 49 yo | Male | African American | Unknown |
| 55 yo | Female | Caucasian | Unknown |
| 28 yo | Female | Hispanic | Unknown |
| 49 yo | Male | Caucasian | Unknown |
| 61 yo | Female | Caucasian | Unknown |
| 56 yo | Female | Caucasian | Unknown |
| 29 yo | Male | Caucasian | Unknown |
| 37 yo | Female | Caucasian | Unknown |
| 55 yo | Female | Caucasian | CNS brain tumor |
| 47 yo | Male | Asian | Anoxia and stroke |
| 49 yo | Female | Caucasian | MVA trauma to head |
| 26 yo | Female | Caucasian | Asphyxiation |
| 45 yo | Male | Caucasian | Anoxia |
| 31 yo | Male | Caucasian | Intracerebral hemorrhage |
| 4 yo | Female | Unknown | Unknown |

The three highest-expanding donors exhibited uniform, cuboidal morphology across both passage 0 (P0) and passage 1 (P1) (Fig. S3A), achieving a cumulative fold expansion exceeding 2,000 per donor by P1, with a mean fold expansion of 4,373 across all donors (Fig. 2C). We scaled the expansion process from 75 cm² flasks to multilayer cell stacks, maintaining consistent yield and viability (Fig. S3B). We routinely generated over 1 billion cells per batch using 10-layer CellStacks, sufficient for preclinical studies, with uniform fold expansion (CV = 15.4%) and cell doublings (CV = 3.9%) across batches (Fig. S3C). Importantly, genomic stability was maintained throughout the expansion process, as the expanded cells retained normal karyotypes (Fig. S3D).

### Expanded human hepatocytes (EHHs) retain their hepatic identity but transiently reduce key functions during expansion

We confirmed that expanded human hepatocytes (EHHs) maintained their hepatic identity and lineage commitment through immunofluorescent staining, showing co-expression of key markers like HNF4α and Albumin in over 99% of cells (Fig. 2D). Gene expression analysis confirmed no significant differences (p > 0.05) in the levels of *ALB*, *CYP3A4*, *HNF4A*, and *SERPINA1* between PHH and EHH or EHH+ cells (Fig. 2E). EHHs upregulated key regenerative pathways, including Wnt, Notch, and HGF/EGF signaling (Fig. S4A), without reverting to a fetal-like state. This was confirmed by negligible *AFP* gene expression in EHHs across multiple donors, as compared to iPSC-derived hepatocytes (iHeps) and fetal hepatocytes (Fig. S4B).

During expansion, EHHs temporarily downregulated critical hepatic functions, as reflected in changes to gene expression, protein levels, and functional activity (Fig. 2Fi, Fig. 2Fii, Fig. 2G respectively). These were largely restored by maturing EHHs in vitro with a growth-factor free, nutrient-rich media formulation over four days. This process produced EHH+ cells with significantly enhanced performance across key axes such as A1AT and Albumin protein secretion (p<1E-7 and p<1E-2, respectively), and CYP3A4 drug metabolism (p<1E-4) (Fig. 2G, Fig. S3E). However, some genes, like *OTC*, did not fully recover (Fig. 2F), resulting in ureagenesis levels that were only 20% of PHH controls (Fig. 2G). To quantitatively benchmark transcriptional profile of expanded hepatocytes, we analyzed their bulk RNA-seq profiles using platform-agnostic CellNet (PACNet), a classification algorithm trained on healthy human samples (21). Reflecting their transient regenerative state, EHHs achieved a mean liver classification score of 0.36. This score increased significantly to 0.7 at the EHH+ stage, approaching the PHH benchmark score of 0.9 (Fig. S4C).

We performed single-cell RNA sequencing (scRNA-seq) on PHH, EHH, and EHH+ cells to evaluate cell-to-cell heterogeneity. PHHs exhibited a highly homogeneous population. However, during expansion at the EHH stage, we identified two distinct subpopulations: one comprised of proliferative, immature cells with high expression of proliferative genes (e.g., *PRC1*, *TOP2A*) and low expression of hepatic genes (e.g., *ALB*, *FGA*, *SERPINA1*, *HNF4A*), and another consisting of less proliferative, mature cells with low expression of proliferative genes and high expression of hepatic genes (Fig. S5). Notably, after in vitro maturation, the heterogeneity resolved from a silhouette score of 0.53 in EHH to 0.25 in EHH+, which was comparable to the PHH silhouette score of 0.34, indicating that the EHH+ cells matured to a more homogenous population comparable to PHH cells (Fig. 2H).

To assess their potential for genetic manipulation, we evaluated the editing efficiency and viability of EHHs across multiple editing modalities. Unlike PHHs, which achieve a maximum of approximately 50% indel efficiency and 60% viability after electroporation (22), EHHs demonstrated higher amenability to gene editing. EHHs achieved >90% viability post-electroporation, >80% efficiency in lentiviral-mediated overexpression of enhanced green fluorescent protein (eGFP), and >90% efficiency in CRISPR-Cas9 mediated knockouts of key human leukocyte antigen (HLA) markers, including β-2 microglobulin (B2M) and class II major histocompatibility complex transactivator (CIITA) (Fig. S6).

### EHHs successfully engraft and repopulate the liver in a chronic liver injury model, leading to improved survival

We tested the engraftment and liver repopulation potential of EHH and EHH+ cells using the *Fah^-/-^/Rag2^-/-^/Il2rg^-/-^*(FRG) model, which has been shown to effectively distinguish between mature and immature hepatocytes (23–27). After injecting 6.5E5 cells into the spleen, we observed migration to the liver and repopulation during 2-(2-nitro-4-trifluoromethylbenzoyl)-1,3-cyclohexanedione (NTBC) ON/OFF cycles (Fig. 3A). By four months post-transplant, EHH and EHH+ cells demonstrated robust integration and functionality, producing an average of 1.4 ± 1.1 mg/mL and 2.2 ± 1.2 mg/mL of human albumin, respectively, compared to 4 ± 0.9 mg/mL from PHHs and significantly lower levels from ‘commercial’ cells (0.3 ± 0.6 mg/mL) and iHeps (0 mg/mL) (Fig. 3B). In this experiment, the ‘commercial’ condition consisted of EHHs cultured in a commercially available media formulation (Takara Bio) that led to poor in vivo engraftment of the cells. Consistent with plasma biomarkers (Fig. 3B and Fig. S7B), histological staining for the human FAH enzyme (FAH-stained fraction: 39.2 ± 5.6% in PHH, 30.8 ± 13.2% in EHH, 31.4 ± 6.9% in EHH+) (Fig. 3C) and quantification of human *Alu* sequences (human DNA/organ: 110.5 ± 37.9 μg in PHH, 20.1 ± 13.0 μg in EHH, 34.2 ± 28.2 μg in EHH+) (Fig. S7D) in mouse liver tissue confirmed human cell repopulation in the EHH and EHH+ groups. Additionally, cells exhibiting higher expression of chemokines and cytokines linked to innate immune signaling showed reduced engraftment efficiency. Notably, EHHs, EHH+ cells, and PHHs displayed lower expression levels of these immune-related molecules, correlating with their improved engraftment outcomes (Fig. 3Dii).

**Figure 3:**
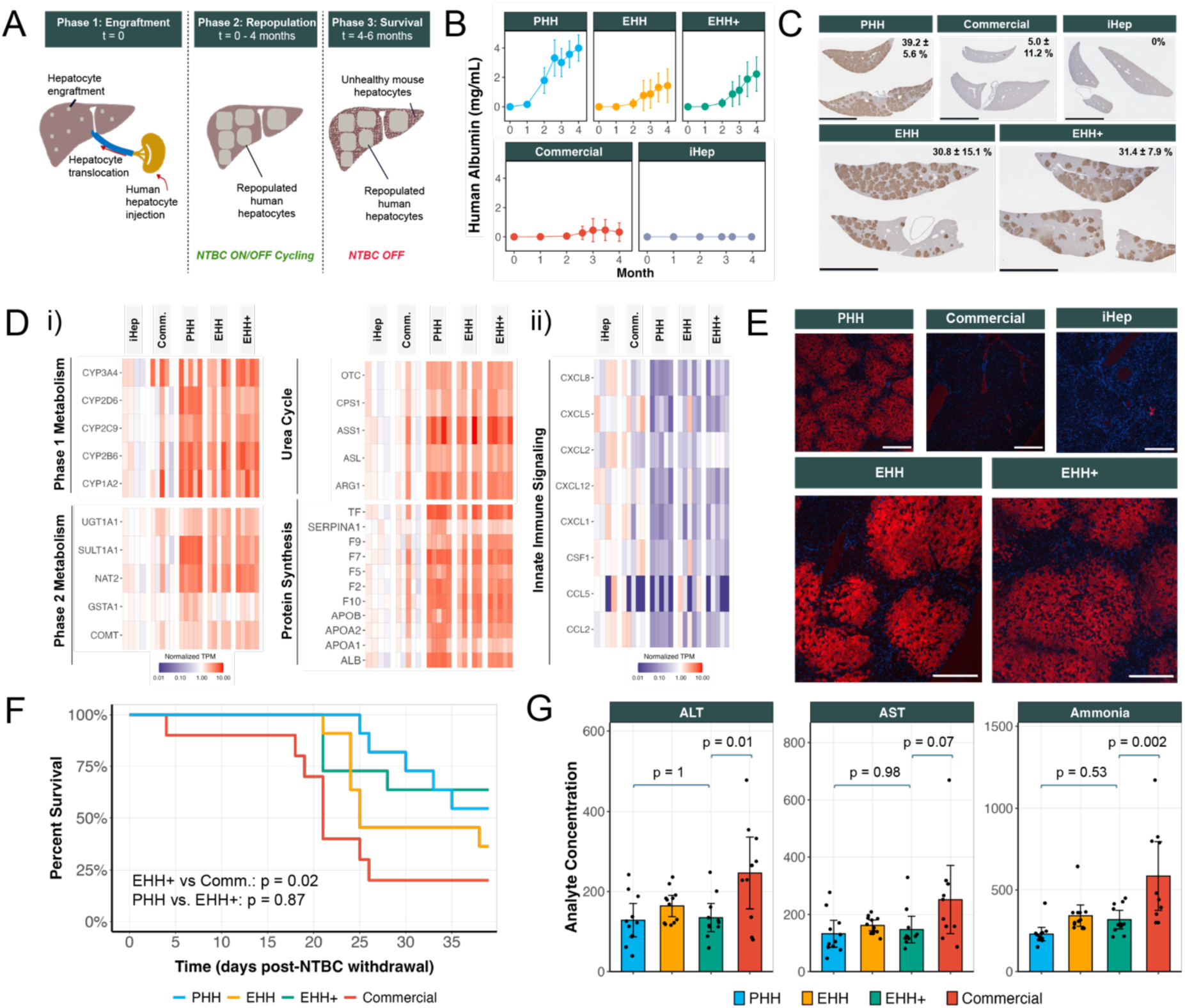
Expanded hepatocytes effectively engraft, repopulate, and rescue liver failure in a hereditary tyrosinemia model. (A) Schematic illustrating the timeline of various phases post-transplantation in FRG mice: Phase 1 (Engraftment, t = 0), Phase 2 (Repopulation, t= 0 - 4 months) where NTBC is cycled, and Phase 3 (Survival, t = 4 - 6 months) where NTBC is withdrawn. (B) Human albumin levels in mouse plasma (mean ± 95% CI) demonstrate successful engraftment and repopulation in PHH, EHH, and EHH+ groups, with no engraftment observed in iPSC-derived hepatocytes and ‘commercial’ cells. (C) Liver explant staining with human fumarylacetoacetate hydrolase (FAH) confirms engraftment patterns consistent with human albumin levels (scale bar = 5 mm). Percentages within each micrograph indicate repopulation rates, quantified from FAH-stained sections (mean ± 95% CI). (D) Normalized expression levels of (i) key phase I and II drug metabolism genes, urea cycle genes, and synthetic pathway genes reveal significant upregulation in cells capable of engraftment and repopulation compared to controls. (ii) In contrast, chemokines and cytokines involved in innate immune signaling are downregulated in engrafting cell populations, suggesting a modulatory role of the innate immune system in facilitating engraftment. Each column represents an individual animal, with data derived from liver explants collected at endpoint (4-6 months). All TPM values are normalized to the mean for the corresponding gene in the iHep group. (E) Representative images of immunofluorescent staining for arginase 1 (ARG1, red) and DAPI (blue) in FRG mouse livers at endpoint show a strong ARG1 signal in engrafted cells compared to iHeps and ‘commercial’ cells (scale bar = 300 μm). (F) Kaplan-Meier survival curves showing enhanced survival rates in animals receiving PHH and EHH+ cells compared to ‘commercial’ cells. The p-values are calculated using the log-rank test. (G) Endpoint liver function tests (ALT, AST) and plasma ammonia levels indicate reduced liver injury in PHH, EHH, and EHH+ groups compared to ‘commercial’ group (mean ± 95% CI). The p-values are calculated using a one-way ANOVA followed by Tukey’s HSD post hoc test for pairwise comparisons. Units for ALT and AST are U/L, while the units for ammonia are μM.

Prior to transplantation, EHH and EHH+ cells exhibited distinct gene expression profiles compared to PHH cells, as demonstrated by UMAP clustering and higher Euclidean distances (p = 1.1E-5; Fig. S7E). However, after four months in vivo in FRG mice, these differences diminished significantly and did not show a significant difference in Euclidean distance (p = 0.35; Fig. S7E). This suggests that the in vivo microenvironment plays a critical role in driving the maturation of EHH and EHH+ cells to closely resemble PHH cells. Engrafted cells in vivo (PHHs, EHHs, and EHH+ cells) showed significant upregulation of genes involved in phase 1 and phase 2 metabolism, protein synthesis, and the urea cycle compared to non-engrafted cells (iHeps and ‘commercial’) (Fig. 3Di). All gene sets demonstrated p-values below 0.01 when comparing iHeps to PHH, EHH, or EHH+ cells (Fig. 3Di). We corroborated this mature phenotype in engrafted cells via high expression of human ARG1 in FRG mouse livers at the endpoint (Fig. 3E).

Finally, to assess therapeutic potential, we performed a survival challenge on the FRG mice after four months by completely withdrawing NTBC (nitisinone), a drug that prevents liver toxicity in this model (Fig. 3A). During the NTBC OFF phase, EHH+ cells significantly improved survival compared to the ‘commercial’ condition (p = 0.02), matching survival fractions observed with PHHs (p = 0.87) (Fig. 3F). Liver injury markers (ALT, AST) and ammonia levels were also significantly lower in the PHH and EHH+ groups compared to the ‘commercial’ condition (ALT: 128 ± 42 U/L for PHH, 135 ± 35 U/L for EHH+ vs. 246 ± 89 U/L for Commercial; ammonia: 229 ± 42 μM for PHH, 318 ± 57 μM for EHH+ vs. 584 ± 210 μM for Commercial) (Fig. 3G), further highlighting their therapeutic potential.

### EHHs can be aggregated with fibroblasts in a controlled manner to create a stable, cryopreserved drug product

To overcome the low engraftment efficiencies commonly observed with single-cell hepatocyte injections in clinical settings, we aggregated EHHs with stromal support cells, specifically normal human dermal fibroblasts (NHDFs), to create 3D constructs called “seeds” (Fig. 4A). Using microwell plates or suspension culture systems, we generated seeds with core-shell structures, with hepatocytes forming the core and NHDFs surrounding at the periphery (Fig. 4B). A rotation speed of 36 RPM can be used to generate 200 µm aggregates, while increasing the speed to 80 RPM reduces the aggregate diameter to 100 µm (Fig. 4Ci). Notably, the aggregate composition remains consistent and can be controlled by the input cell ratio, irrespective of the rotation speed (Fig. 4Cii). Hepatocytes within the seeds preserved their identity, as demonstrated by the co-expression of HNF4α and Albumin (Fig. S8A). They maintained high viability (>70%) for over 24 hours (Fig. S8B) and exhibited significantly higher viability than PHH single cells (EHH seeds: 87.5% vs. PHH cells: 74.4%; p = 1.1E-5) (Fig. S8C).

**Figure 4:**
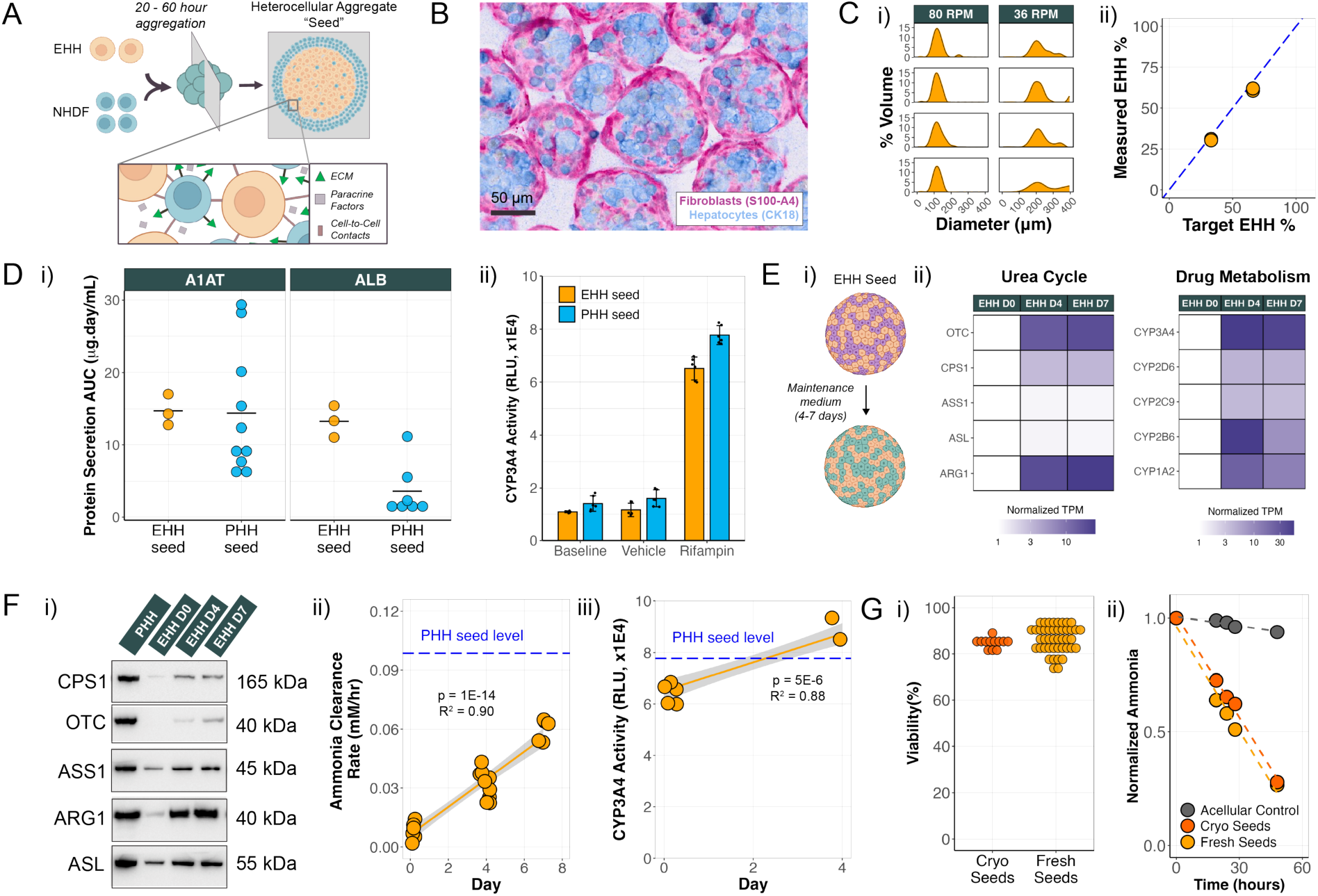
Expanded hepatocyte aggregation into heterocellular constructs leads to a cryopreserved drug product with enhanced functional maturation. (A) Schematic depicting the assembly of hepatocyte-fibroblast seed constructs, supported by extracellular matrix (ECM), paracrine, and cell-cell interactions. (B) Histological staining (S100-A4 for fibroblasts, CK18 for hepatocytes) highlights the integration of the two cell types within a core-shell seed structure. (C) (i) Assessment of seed volumetric distribution in bioreactors across two different RPMs, demonstrating a scalable production process with size control (each row represents an individual batch). (ii) Measurement of EHH cell fraction using short-tandem repeat (STR) analysis, showcasing the ability to achieve multiple target cell ratios. (D) (i) A1AT, albumin production levels (n = 3 donors for EHH seeds and n = 7-10 donors for PHH seeds, and (ii) CYP3A4 induction in EHH seeds versus PHH seed control groups, indicating comparable function to primary cells (n = 1 donor, n = 6 technical replicates). (E) Maturation of EHHs in seed format is achieved by culturing in maintenance media for 4 - 7 days, as demonstrated by enhanced expression profiles of key genes involved in the urea cycle and drug metabolism. All TPM values are normalized to the EHH D0 group for each corresponding gene. (F) Maturation in seed format over time is confirmed by (i) elevated protein expression of key urea cycle enzymes, measured by Western blotting, (ii) enhanced ammonia clearance rate (n = 1 donor, n = 5-13 technical replicates per timepoint), and (iii) enhanced cytochrome p450 3a4 metabolism post-induction with rifampin (n = 1 donor, n = 6 technical replicates per timepoint), indicating functional improvement. Functional levels for corresponding PHH seeds are indicated by blue dashed lines. (G) Demonstrating successful cryopreservation of seeds, evidenced by comparable (i) cell viabilities (n = 12 batches for cryopreserved seeds and n = 44 batches for fresh seeds) and (ii) ammonia clearance capacity between fresh and cryopreserved seeds (ammonia values are normalized to t = 0 hours for each group).

We compared EHH seeds to PHH seeds and found equivalent or superior albumin and A1AT secretion in vitro (Fig. 4Di), and CYP3A4 activity in EHH seeds showed a 5.5-fold increase upon rifampin induction, reaching 78% of PHH seed levels at baseline and 84% following induction (Fig. 4Dii). To further increase their maturation state, we cultured the seeds in maintenance media for 4-7 days (Fig. 4Ei). This was demonstrated by the upregulation of urea cycle and drug metabolism genes (Fig. 4Eii) and an increase in the levels of urea cycle proteins (Fig. 4Fi). Additionally, protein secretion and metabolic activity improved during this culture period, as evidenced by an 8-fold increase in ammonia metabolism (Fig. 4Fii), an increase in ureagenesis (Fig. S8E), and a 1.4-fold increase in induced CYP3A4 activity (Fig. 4Fiii). The presence of NHDFs was critical as it enhanced functions such as ammonia clearance by 2.8-fold compared to seeds lacking NHDFs (p = 3E-4) (Fig. S8D).

We developed a cryopreservation process to improve seed stability and to support its viability as a commercial drug product, enabling full batch release and testing prior to administration. Post-thaw, the seeds demonstrated recoveries around 90% (Fig. S9A), maintained their pre-cryopreservation size distribution (Fig. S9B), and exhibited comparable viability (fresh: 86.3%, cryopreserved: 85.0%, p = 0.43) and ammonia clearance functionality to fresh seeds (Fig. 4G). Additionally, cryopreserved EHH seeds exhibited protein secretion levels comparable to those of cryopreserved PHH seeds (Fig. S9C).

### Seed configuration enhances engraftment efficiency, which can be further improved through innate immune suppression

To evaluate the engraftment potential of seeds across multiple anatomic sites, we employed the NSG mouse model (Fig. 5A) and compared NHDF-seeded constructs to fibroblast-free seeds and seeds aggregated with bone marrow-derived mesenchymal stromal cells (BM-MSCs) in the kidney capsule. NHDF seeds demonstrated significantly better engraftment compared to fibroblast-free and BM-MSC seeds, as evidenced by elevated plasma human albumin levels (AUC: 258 µg·day/mL for NHDF seeds vs. 48 µg·day/mL for BM-MSC seeds, p < 1E-4) and the presence of hepatocytes on explant confirmed by hematoxylin and eosin (H&E) staining and carbamoyl phosphate synthetase 1 (CPS1) expression (Fig. 5B). Aggregation with fibroblasts enabled successful engraftment across extrahepatic sites, including the spleen, fat pad, and kidney capsule, as evidenced by stable human albumin levels and positive human OTC staining in explanted tissues (Fig. 5C).

**Figure 5:**
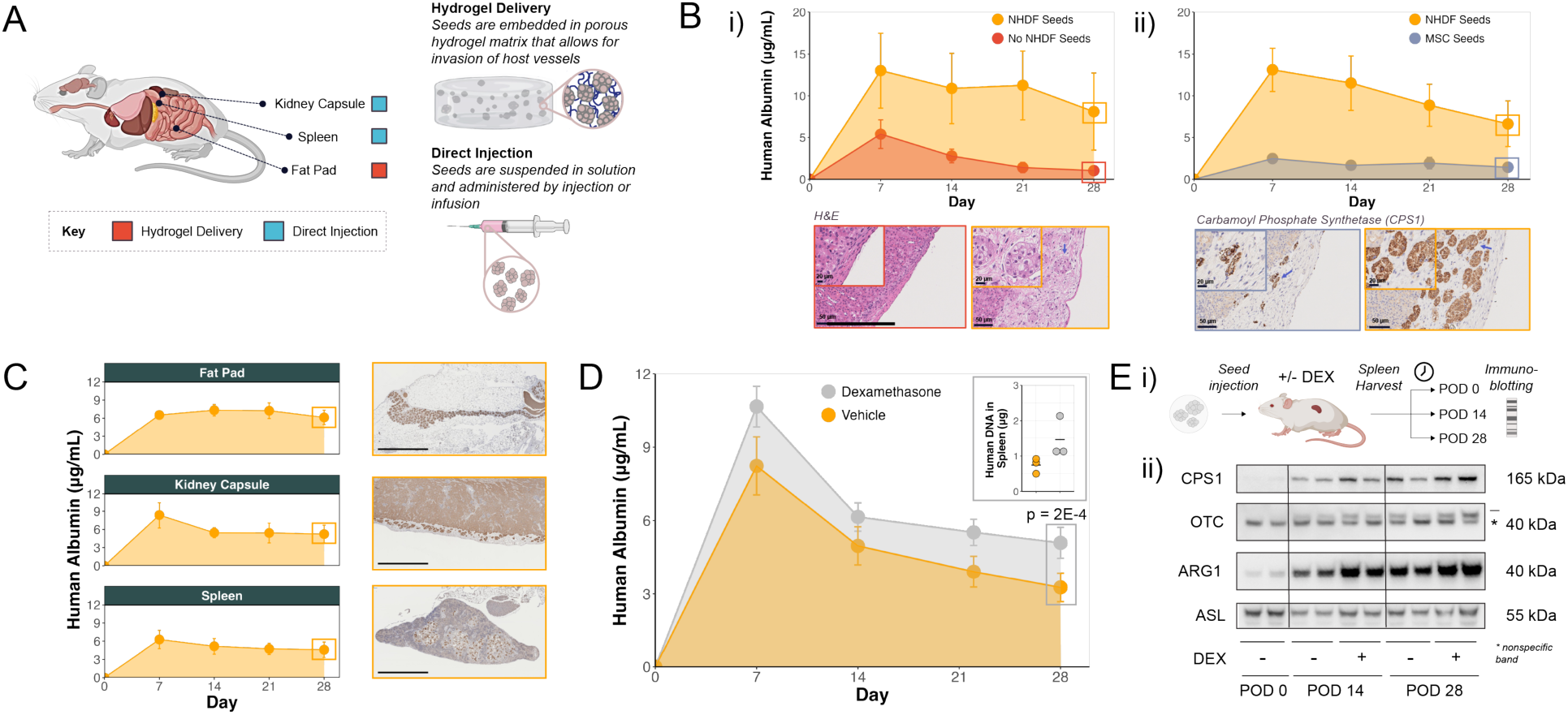
Seeds enable engraftment across various extrahepatic sites and engraftment can be improved by innate immune suppression. (A) Schematic overview of seed implantation in various anatomical sites (fat pad, kidney capsule, spleen) using both hydrogel and direct injection methods. (B) Heterocellular aggregation of EHHs with dermal fibroblasts significantly enhances engraftment within the kidney capsule, as evidenced by increased plasma albumin levels (mean ± 95% CI) and histological comparisons showing reduced engraftment in (i) the absence of dermal fibroblasts or (ii) when using bone marrow-derived mesenchymal stromal cells (BM-MSCs). (C) Time-course of human albumin levels (mean ± 95% CI) and immunohistochemical analysis of ornithine transcarbamylase (OTC) across implantation sites (fat pad, kidney capsule, spleen) shows that seed constructs support stable engraftment and sustained hepatocyte activity across various extrahepatic locations (scale bar = 1 mm). (D) Innate immune suppression with glucocorticoid (dexamethasone) administration enhances engraftment within the spleen, evidenced by increased human albumin levels (mean ± 95% CI) and a higher abundance of human DNA in mouse spleen samples at POD28 (inset). The p-value is calculated using a two-sample t-test to compare albumin levels at POD28. (E) (i) Schematic illustrating in vivo assessment of functional maturation. (ii) Western blot analysis of human liver-specific proteins in the spleen indicates progressive functional maturation of EHH seeds in vivo over time, both with and without glucocorticoid administration. All antibodies were validated to ensure they did not cross-react with mouse spleen tissue.

Building on insights from the FRG model, where poor engraftment was associated with high expression of innate immune signaling molecules (Fig. 3Dii), we hypothesized that further suppressing the innate immune response could improve engraftment outcomes. To test this hypothesis, we administered the glucocorticoid dexamethasone in the NSG mouse model. This treatment significantly improved seed engraftment in the spleen, as shown by elevated human albumin levels (AUC: 132 µg·day/mL for vehicle vs. 175 µg·day/mL for Dexamethasone, p < 0.01, Fig. 5D), a 1.9-fold increase in human *Alu* copies in explanted tissues (Fig. 5D), and notably larger seed beds grossly detected in H&E-stained explants (Fig. S10A). Additionally, we observed rapid in vivo maturation of the seeds, which began as early as post-operative day (POD) 2 (Fig. S10B). By POD14, the maturation process stabilized, marked by significant upregulation of urea cycle enzymes (Fig. 5E).

### Seeds can be localized to the spleen via splenic artery delivery in large animals

We tested a clinically relevant delivery route by administering seeds into the splenic artery of Göttingen minipigs using a direct catheterization procedure and controlled infusion rate (Fig. S11A). The procedure was well tolerated, all animals fully recovered, and no adverse events were reported. Biopsy analysis using immunohistochemistry for human CK18 (Fig. S11B) and quantification of human *Alu* sequences (Fig. S11C) revealed that the seeds predominantly localized to the spleen (mean: 4,623 μg human DNA in the spleen vs. 170 μg in the liver, with no detection in off-target organs). In contrast, single-cell hepatocyte transplants typically migrate to the liver and can have the potential for a wider off-target distribution to other organ systems resulting in safety concerns (6). These findings highlight the potential for the seeds to localize and engraft in desired extrahepatic sites following intraarterial delivery, which may translate to therapeutic and safety benefits.

## Discussion

Clinical application of hepatocyte-based therapies is limited by the lack of a scalable source of functional hepatocytes and poor engraftment. To address these challenges, we developed a cGMP-compliant, scalable process for expanding primary hepatocytes while preserving their functionality and improving engraftment through heterocellular aggregation with supportive cells, resulting in a stable, cryopreserved therapeutic product.

We demonstrate that our proprietary media formulation enables the direct expansion of PHHs without inducing dedifferentiation or requiring subsequent re-differentiation. This process mirrors the regenerative response observed following liver resection, where adult hepatocytes transiently downregulate liver-specific functions to allocate bioenergetic resources toward cell cycle entry and progression (28). Importantly, this temporary downregulation of function is quickly restored across various contexts, including 2D cultures (Fig. 2G), heterocellular aggregates in suspension (Fig. 4F), and directly in vivo (Fig. 3D, Fig. S7E, Fig. 5E). Furthermore, we show that PHHs can be rapidly expanded using entirely xenogeneic-free formulations, an advancement over prior methods that require several months of prolonged culture with xenogeneic components such as fetal bovine serum (FBS) and rat-tail collagen (24), or rely on 3D culture systems using mouse sarcoma-derived Matrigel (29). Our protocol achieves comparable expansion within approximately three weeks, enabling the production of large, clinically relevant cell batches for therapeutic applications. Our findings also suggest that specific subpopulations may play a key role in driving hepatocyte expansion. Notably, *PROCR* is expressed in 76.3% cells during the expansion phase but is limited to a small subset in parent PHHs and mature EHH+ cells (Fig. S5D). Originally identified in pancreatic literature (30), *PROCR* highlights potentially conserved biology across endodermal lineages and opens opportunities to refine methods by isolating and targeting these subpopulations to enhance proliferation.

EHHs offer significant advantages over iPSC-derived hepatocytes (iHeps), which are often proposed as an alternative source of hepatocytes. While iHeps are highly scalable and exhibit robust albumin expression, they struggle to perform advanced liver functions that require complex metabolic inputs, such as ammonia and drug metabolism (31–34). In contrast, EHH cells and seeds retain a more mature hepatic phenotype, with high CYP3A4 activity (Fig. 4D) and urea cycle function (Fig. 4F), and do not express fetal markers like AFP, which are much higher in iHeps (Fig. S4B). iHeps effectively display an epigenetic and transcriptomic profile that aligns more closely with fetal hepatocytes than adult PHHs, including incomplete silencing of pluripotency-related gene (7). Additionally, iHeps exhibit poor in vivo engraftment, as shown in the FRG mouse model, where EHH+ cells achieved significantly better engraftment and functional integration (Fig. 3). For iHeps to become a viable source of hepatocytes for clinical applications, differentiation protocols must overcome their fetal-like stage and advance their functional maturity. Until then, their therapeutic potential remains limited, further emphasizing the importance of direct PHH expansion as a scalable and functional source of hepatocytes.

A key challenge in the field is genetically engineering PHHs with high efficiency and viability. Once isolated from their native liver microenvironment, PHHs are highly susceptible to anoikis and in vitro stress, often losing liver-specific functions and viability (35–37). Genetic manipulation techniques, such as electroporation, viral transduction, or chemical exposure, can further exacerbate this sensitivity, frequently leading to cell death. As terminally differentiated, non-dividing cells, PHHs present a hurdle for genetic editing methods like CRISPR-Cas9, which depend on cellular repair pathways typically active during cell division (e.g., homology-directed repair or non-homologous end joining) (38). To address this limitation, we genetically edited EHHs during their proliferative stage and observed remarkably high editing efficiencies and high viability post-editing (≥ 90%, Fig. S6), significantly surpassing those reported in the literature for PHHs (22). This finding underscores the potential of using EHHs as a platform for genetic modifications, paving the way for novel therapeutic applications that leverage engineered hepatocytes.

Additionally, we demonstrated that aggregation with stromal cells results in higher function (Fig. S8D), improved stability (Fig. S8B), a more localized biodistribution profile (Fig. S11C), and enhanced engraftment outside the liver (Fig. 5B). In contrast to 2D culture, where EHHs did not recover OTC expression post-maturation (Fig. 2Fii) and achieved only 20% of the ureagenesis levels observed in PHHs (Fig. 2G), the seed configuration further improved maturation, evidenced by increased OTC expression (Fig. 4Fi) and achieving 60% of PHH ammonia clearance levels (Fig. 4Fii). We hypothesize that these benefits come from a combination of paracrine factors (19, 20), extracellular matrix interactions (36, 39, 40), and cell-cell signaling (41–43) provided by the stromal cells, which together create a supportive microenvironment for hepatocyte stabilization. If the key factors driving these effects can be identified, it may be possible to replicate them synthetically, eliminating the need for stromal cells. Ectopic engraftment is particularly valuable for chronic liver disease, where fibrosis and inflammation often make the liver an unsuitable site for cell therapy (17). Future studies should evaluate the therapeutic efficacy of seeds in preclinical models of fibrotic liver disease. These studies could also help define the minimum cell mass necessary at an ectopic site to achieve therapeutic efficacy.

Finally, the role of innate immunity in hepatocyte engraftment warrants further exploration. Our experiments demonstrated significant improvement in engraftment through suppression of the innate immune response (Fig. 5D), but additional work is needed to deepen our understanding of this mechanism. Future studies could leverage knockout mouse models lacking specific immune system compartments to dissect the contributions of various innate immune effectors. Genetically edited EHHs designed to evade or modulate immune responses also offer a promising avenue for improving engraftment outcomes.

Overall, we provide evidence that the scalable expansion of PHHs and aggregation with stromal cells represents a step forward in overcoming barriers that have historically hindered the translation of hepatocyte-based therapies into clinical use.

## Materials and Methods

### Isolation of hepatocytes from human livers

Hepatocytes were isolated from human cadaveric livers using similar protocols as described in the literature (44) and cryopreserved in a DMSO-based cryostorage medium utilizing xenogeneic-free reagents throughout the process.

### Expansion of primary human hepatocytes

Cryopreserved PHH were thawed, diluted with PHH thaw media, and concentrated using centrifugation. After the cells were centrifuged, they were resuspended in plating medium and counted. Based on the viable cell fraction, the cells were seeded into tissue culture vessels coated with xenogeneic-free extracellular matrix, marking Day 0 of the process. The cells were plated and allowed to adhere overnight under normoxic (21% O₂) cell culture conditions at 37°C and 5% CO₂. After overnight culture, the cells were transferred to hypoxic cell culture conditions. The cells were fed every other day using a proprietary xenogeneic-free media (Satellite Bio) formulated to support the expansion of PHH. After the cells reached confluence, the cells were lifted with Accutase (Innovative Cell Technologies) and concentrated.

### Expansion of normal human dermal fibroblasts (NHDFs)

A vial of NHDF was thawed and seeded into a 2-layer cell stack at 1.5-2E3 viable cells/cm². The cells were cultured in growth xenogeneic-free growth media in a 37°C, 5% CO₂ incubator until confluent, with media changes every two to three days. The fibroblasts were then passaged using TrypLE Select (Gibco), centrifuged at 340×g for 10 minutes, and seeded into 10-layer cell stacks (Corning) at 1.5-2E3 viable cells/cm². The cells were expanded again under the same growth conditions until confluent, with media changes every two to three days. The cells were then harvested using TrypLE Select, centrifuged at 340×g for 10 minutes, and cryopreserved at 5E7 viable cells/mL in CryoStor® 10 (BioLife Solutions).

### Expansion of bone-marrow derived mesenchymal stromal cells (BM-MSCs)

BM-MSC vials were cultured per manufacturer’s instructions (Rooster Bio) and cryopreserved at 2-5E6 viable cells/mL in CryoStor® 5.

### Cryopreservation of PHHs and expanded human hepatocytes (EHHs)

Cells were resuspended in 2x cryostorage diluent medium. The cells were counted, and the number of cryopreservation vessels was determined based on the viable cell count. The cells were then resuspended in 2x cryopreservation media (Satellite Bio), aliquoted into cryovessels, and placed in a Controlled Rate Freezer (CRF) where they were cryopreserved using a custom profile. After the CRF run was completed, the cells were moved to long-term storage in the vapor phase of liquid nitrogen.

### 2D maturation of EHHs

After the expansion phase, EHHs were matured (EHH+) for an additional 4 days in a normoxic incubator (37°C, 5% CO₂, 20% O₂), with a media exchange on day 2. The EHH+ cells were cultured in growth-factor free, nutrient-rich hepatic maturation media (Satellite Bio). The complete maturation media were prepared fresh and pH balanced prior to use on the cells.

### G-banding karyotyping of EHHs

Cells were incubated with colcemid (Life Technologies) at 37°C, 5% CO₂ for approximately 80 minutes. After the incubation period, the cells were harvested, treated with potassium chloride, and then fixed with methanol and glacial acetic acid. Chromosomes were then assessed for abnormalities via G-banding. G-banding was performed by Cell Line Genetics.

### Cell culture of control lines

PHHs cultured using LifeNet Health media: Wherever necessary as positive controls, PHHs were thawed using hepatocyte thaw medium (Gibco) and plated on Cultrex rat collagen 1 (Bio-techne) coated culture vessels at a density of 2 mL per well of a 6-well plate at 0.75E6-1.5E6 cells/mL in hepatocyte plating media (HHPM, LifeNet Health®, MED-HHPM) for 4 hours. Following 4 hours of plating, the media were aspirated, and fresh, complete hepatocyte culture media (HHCM, LifeNet Health®, MED-HHCM) were added. Cells were cultured for a maximum of four days, with a media change on day 2, or harvested after one day in culture.

EHHs cultured using Takara media: EHHs were thawed and plated at a density of 67E3 cells/cm^2^. Cellartis® Power™ Primary HEP Medium (Takara) was added after the 4-hour plating period and for the media change on day 2.

iPSC-Derived Hepatocytes (iHEP): When required, iHEPs (FujiFilm Cellular Dynamics) were cultured using CELLCOAT® collagen type 1 pre-coated plates (Greiner Bio-One) following the manufacturer’s guide. Cells were thawed and plated at a seeding density of 3.0E5/cm² using plating media comprising RPMI 1640 Medium (Thermo Fisher) supplemented with iCell hepatocyte 2.0 medium supplement (FujiFilm Cellular Dynamics), 1X B-27 (Thermo Fisher), 20 ng/mL rh-Oncostatin-M (Bio-techne), 0.1 µM Dexamethasone (MP Biomedicals), and 25 µg/mL gentamicin (Thermo Fisher). After 4 hours, 100% of the media were exchanged with fresh plating media. Cells were cultured for 5 days in plating media, with a media exchange every day. On day 5, cells were transitioned to maintenance media, which were the same as plating media without Oncostatin-M. Cells were cultured for an additional 48 hours and subsequently harvested.

### Lentiviral-mediated overexpression in cells

PHHs were thawed and plated as previously described for expansion. A lentiviral construct containing the transgene of interest under the SFFV promoter was cloned and packaged into lentiviral particles by VectorBuilder. Cells were expanded for 7 days before being transduced with lentiviral particles at specific multiplicity of infections (MOI) for the delivery of the transgene of interest (eGFP or NLuc). Twenty-four hours post-transduction, fresh expansion media was added, and the cells were cultured for 2 additional days. On day 10 of culture, antibiotic selection was initiated using expansion media containing 4 µg/mL Puromycin (ThermoFisher) for eGFP or 100 µg/mL Hygromycin B (ThermoFisher) for NLuc, continuing for 7 days. On day 17 of culture, cells were assessed for eGFP expression using Cytation 10 or for NLuc activity using the Nano-Glo luciferase assay (Promega).

### CRISPR-Cas9 mediated knockout in cells

For CRISPR-Cas9 knockout of specific target genes, sgRNAs for β-2-microglobulin (B2M, Synthego, CAGCCCAAGAUAGUUAAGUG) and class II MHC transactivator (CIITA, Synthego, CAUCGCUGUUAAGAAGCUCC) were complexed with Cas9s (Synthego) to form ribonucleoproteins (RNPs). Next, thawed EHHs were mixed with the RNPs and electroporated using a Neon™ NXT electroporator (ThermoFisher Scientific). After electroporation, the cells were placed in a T-75 flask with expansion media supplemented with a small molecule cocktail that aids cell viability and functionality post genetic editing (45). The edited cells were cultured for an additional 11 days, with a complete media change every other day.

At the end of the culture, the cells were counted, and their genomic DNA (gDNA) was extracted using the Maxwell RSC instrument (Promega). sgRNA and target-specific PCR primers were designed and used to amplify the edited region in the extracted gDNA using a thermocycler (Fisher). Amplicon samples were sent to a third-party vendor, GeneWiz, for Sanger sequencing, and the resulting .ab1 files were analyzed using Synthego’s Inference of CRISPR Edits (ICE) tool to determine the editing efficiency achieved.

### Aggregation of EHHs to form seeds

Aggrewells™ 400 6-well plates (Stemcell Technologies™) were utilized for small scale aggregations. Briefly, the wells were coated with anti-adhesion solution (Stemcell Technologies) and subsequently washed with PBS (Gibco™) following the manufacturer’s protocol. Next, a cell seeding solution of NHDFs and EHHs, each at appropriate concentrations, was prepared and 2mL of this was added to each well. The plate was spun at 60g for 6min and placed in a normoxic incubator for 24hrs. Following aggregation, seeds were either harvested or further cultured dictated by downstream needs.

For larger scale aggregations, 0.1L Vertical Wheel Bioreactors (VWB, PBS Biotech) were inoculated with EHHs and NHDFs to a total volume of 80–100 mL in aggregation media. The reactors were placed on bases set to 36 -80 RPM in an incubator (37°C, 5% CO₂). Shortly after starting agitation, a sample of cell suspension was taken from each bioreactor for analysis of metabolites and measurement of cell count and viability. After 20-24 hours, the bioreactors were prepared for the harvest of the aggregates. A sample of cell/aggregate suspension was collected from each bioreactor for analysis of metabolites. One to five 0.1L VWBs were harvested into a 1L transfer bag. The seeds were then washed using a commercial cell washer to remove single-cell debris and exchange media from aggregation to wash media. After the wash step, aggregates collected from the bioreactor were sampled and held at 4°C until use or cryopreserved.

### Quantification of seed size and volume

Seed size and volume were measured by first aliquoting seeds into a custom 10 mm glass imaging chamber, followed by capturing a phase contrast image of the whole field using the EVOS M7000 imaging system. Seeds were then segmented from the background using Otsu’s threshold (46), and individual seeds were identified and counted based on connected, thresholded pixels. Seed-to-seed contact was detected by applying an area-to-perimeter ratio cutoff, and any particle exceeding this cutoff was subjected to watershed segmentation to discretize individual seeds. An equivalent diameter was calculated, assuming the thresholded pixel area of each seed represented a circle. Finally, the volume of each seed was calculated by using the equivalent diameter as the diameter of a sphere.

### Short-tandem repeat (STR) analysis of seeds

DNA was extracted and purified using the Maxwell RSC instrument and consumables (Promega). The total amount of extracted DNA was quantified using UV-Vis and then analyzed using the Geneprint 24 system (Promega) for its Short Tandem Repeat profile. The ratio of fibroblasts to hepatocytes was determined using the ChimerMarker software and the unique donor fingerprints. The total cell concentration was calculated using the known genomic copies per cell for the hepatocytes and fibroblasts.

### Immunocytochemistry (ICC) of cells and seeds

Cells and/or 3D seeds were stained for specific target proteins via routine immunofluorescence protocols. Briefly, the cell/3D seeds were washed with PBS (Gibco™), fixed in a paraformaldehyde (PFA) fixative solution (ThermoFisher), and subsequently permeabilized with 0.4% TritonX-100 (Sigma-Aldrich). Following permeabilization, cells/seeds were washed and blocked in either bovine serum albumin (BSA, Sigma-Aldrich) or normal donkey serum containing solutions. After blocking, respective primary antibodies were added, and the cells/seeds were incubated overnight at 4°C. After overnight incubation, cells/seeds were washed with 0.1% Tween 20 (Sigma-Aldrich, P9416) or just PBS wash buffer and incubated for 1hr, in the dark, at room temperature with the appropriate fluorophore-conjugated secondary antibodies. Cellular nuclei were stained using Hoechst 33342 (Invitrogen, H3570). Finally, cells/seeds were washed three times with a wash buffer and then imaged using a Cytation 10 spinning disk confocal microscope (Agilent Technologies). With regards to the 3D seeds, z-stack fluorescence images were captured with a step size of 5–10 µm intervals using a 20X objective. Images were portrayed as maximum z-stack projections.

Antibodies and dilutions used herein are listed in Table 4. A rabbit IgG isotype (Cell Signaling Technology) and a mouse IgG1 isotype (Cedarlane) were used as isotype controls for staining in equal amounts to their respective antibodies.

**Table 3:** Demographic information for all stromal cell donors.

| **Cell Type** | **Age** | **Sex** | **Race** |
| --- | --- | --- | --- |
| NHDF | Neonate | Male | Unknown |
| BM-MSC | 20 yo | Female | Unknown |

**Table 4:**
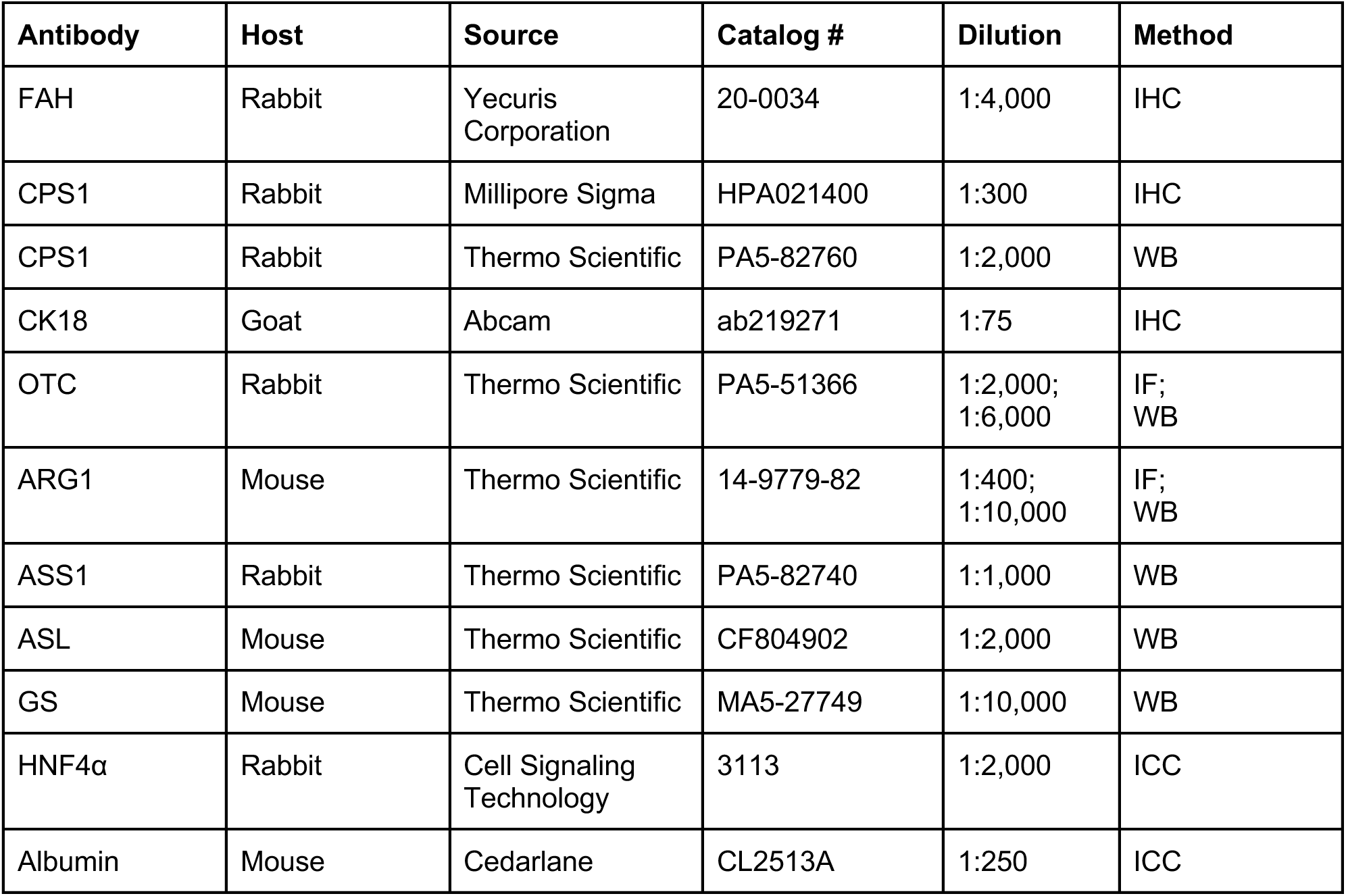
List of antibodies used for Immunohistochemistry (IHC), Immunofluorescence (IF), Immunocytochemistry (ICC), and Western blotting (WB)

| **Antibody** | **Host** | **Source** | **Catalog #** | **Dilution** | **Method** |
| --- | --- | --- | --- | --- | --- |
| FAH | Rabbit | Yecuris Corporation | 20-0034 | 1:4,000 | IHC |
| CPS1 | Rabbit | Millipore Sigma | HPA021400 | 1:300 | IHC |
| CPS1 | Rabbit | Thermo Scientific | PA5-82760 | 1:2,000 | WB |
| CK18 | Goat | Abcam | ab219271 | 1:75 | IHC |
| OTC | Rabbit | Thermo Scientific | PA5-51366 | 1:2,000;<br>1:6,000 | IF;<br>WB |
| ARG1 | Mouse | Thermo Scientific | 14-9779-82 | 1:400;<br>1:10,000 | IF;<br>WB |
| ASS1 | Rabbit | Thermo Scientific | PA5-82740 | 1:1,000 | WB |
| ASL | Mouse | Thermo Scientific | CF804902 | 1:2,000 | WB |
| GS | Mouse | Thermo Scientific | MA5-27749 | 1:10,000 | WB |
| HNF4α | Rabbit | Cell Signaling Technology | 3113 | 1:2,000 | ICC |
| Albumin | Mouse | Cedarlane | CL2513A | 1:250 | ICC |

### Maintenance of seeds in culture

Where necessary, aggregated seeds were cultured in complete human hepatocyte culture media (HHCM, LifeNet Health®, MED-HHCM) in Aggrewells™ 400 6-well plates (Stemcell Technologies™) for either 4 days or 7 days. Wells were coated with 2 mL of anti-adhesion solution (Stemcell Technologies) for 30 minutes according to the manufacturer’s protocol. The wells were subsequently washed three times with 2 mL of PBS (Gibco™) at room temperature.

Once aggregated, 3.6 µL of seed volume was placed in each pre-coated well using a wide-bore, pre-passivated tip. The plates were spun down at 200g for 5 minutes to ensure the seeds settled to the bottom of the microwells. The plates were then placed in a humidified, normoxic incubator. An 80% media change was carefully performed every other day to ensure no seeds were aspirated. If needed, supernatants were saved for additional downstream analyses.

### Cryopreservation of seeds

After washing, seeds were centrifuged at 400×g for 5 minutes, and the pellet was resuspended in pre-cooled cryopreservation media (Satellite Bio) at a concentration of 60 - 150 µL seeds/mL. The suspension was split across cryovials. The seeds were then frozen using a Controlled Rate Freezer (CRF) and transferred to the vapor phase of liquid nitrogen for storage.

### Encapsulation of seeds in fibrin hydrogels

For encapsulation, seeds were resuspended in 157 µL of Fibryga (14 mg/mL). The suspension was rapidly mixed with 157 µL of 2.5 U/L Recothromb before being cast into two pre-passivated 10 mm diameter molds. The grafts were incubated at 37°C for 45 minutes before being removed with a spatula and transferred to a conical flask containing DMEM + 0.5% human serum albumin for transport at 4°C.

### Measurement of cell viability in seeds

Viability was measured using a nuclear stain approach. Seeds were first double-stained with NucFix Red and Hoechst 33342 to identify dead nuclei and all nuclei, respectively. Seeds were then fixed in formalin, aliquoted into a glass imaging chamber, and placed in a Cytation 10 spinning disk confocal microscope. A z-stack of fluorescence images was captured at 20x magnification with 15 µm intervals in the z-plane, using 361 nm excitation / 497 nm emission for Hoechst and 520 nm excitation / 593 nm emission for NucFix.

Each image was contrast maximized and subjected to an unsharp mask, and images exceeding a blur threshold were automatically removed. Individual nuclei were segmented using a neural network, and the number of all nuclei and dead nuclei across the entire stack was summed. Finally, viability was calculated as: (n_all_nuclei – n_dead_nuclei) / n_all_nuclei.

### Measurement of protein secretion from cells

Secreted proteins were quantified by ELISA. Cell culture supernatants were collected at different time points during the various stages of 2D cell and 3D seed culture. Collected supernatants were diluted appropriately to obtain concentrations within the linear range of the standard curve for each analyte tested. Levels of human albumin (Bethyl Laboratory), α1-antitrypsin (Bethyl Laboratory), and Factor-IX (Abcam) were measured according to the respective manufacturers’ protocols.

### Measurement of protein secretion from seeds

Protein secretion was measured in vitro by encapsulating seeds in fibrin gels and changing media every other day for 10 days. At every media change, an ELISA was performed to measure protein concentrations of albumin and A1AT. After three media changes (4 days) to wash out contributions of residual protein from the manufacturing process, a relative protein secretion was calculated as the area under the curve (AUC) of concentrations from day 4 to day 10.

### Measurement of CYP3A4 activity and induction

For the measurement of CYP3A4 metabolism activity, each cell type (EHH or EHH+) was cultured according to the previously described protocols. After each stage, donor-matched PHHs were plated and used as a positive control. Culture media was removed, and Cellartis® Power™ Primary HEP Medium (Takara) was added to the cells and cultured for an additional 72 hours, with media changed every day. Specific wells were designated for baseline controls, vehicle controls (DMSO, Sigma-Aldrich), and 25 µM rifampicin (Sigma-Aldrich)-induced CYP3A4 activity. After 72 hours, CYP3A4 activity levels were measured using a luminescent method with the P450-Glo™ CYP3A4 assay (Promega) following the manufacturer’s instructions.

Briefly, the wells were washed with PBS (Gibco), following which a 3 µM substrate solution was added to each well and incubated at 37°C for 1 hour. Next, equal volumes of supernatant and reconstituted luciferin detection reagent were mixed and incubated further at room temperature. Finally, luminescence was measured using a Tecan plate reader and the induced CYP3A4 activity was calculated by normalizing to the baseline samples and the vehicle samples.

Additionally, the CYP3A4 activity of 3D aggregated seeds was measured following a similar protocol as above, with minor changes. First, seeds were maintained in complete LNH-HHCM (LifeNet Health) media according to previously described maintenance steps. Moreover, the 72-hour CYP induction phase occurred in LNH-HHCM (LifeNet Health) media, with specific wells once again allocated for baseline, DMSO, and 25 µM rifampicin-induced CYP activity. Following the induction, the same P450-Glo assay kit was used to detect and calculate enzyme activity within the 3D seeds.

### Measurement of native and 15N-ureagenesis

Native Ureagenesis: Cell culture supernatants at various timepoints during different stages of cell culture were analyzed for the presence of urea using a QuantiChrom™ urea assay kit (BioAssay Systems, DIUR-100). In accordance with the manufacturer’s instructions, culture media supernatants were assayed directly, in duplicate, using a 96-well plate. Additionally, the provided urea standard was spiked into the respective media used in the experiment to prepare a standard curve for each assay. Optical density (O.D.) measurements were taken at 430 nm or 520 nm, depending on the expected urea concentrations, using a Tecan 200Pro plate reader.

15N-Ureagenesis: Aggregated seeds were cultured in the presence of 15N-labeled ammonia-containing media as described elsewhere for assessing ammonia clearance. Cell culture supernatants were analyzed after the ammonia clearance assay for the production of 15N-urea. Assay media supernatants were collected and sent to NovaBioAssays, LLC (Woburn, MA) for LC-MS-based measurement and quantification of the 15N-labeled heavy urea analytes.

### Measurement of ammonia clearance

Culture media spiked with ammonium chloride was inoculated with seeds and cultured over a 48-hour period. Supernatant samples were collected over two days after the ammonia challenge. The supernatants were then assessed for remaining ammonia levels using the Cedex BioHT (Roche) analyzer with the NH3Bio HT assay. Linear slopes were determined from a zero-order fit to calculate the ammonia clearance rate (mM/hr).

### Measurement of protein expression via Western Blotting

Cells were lysed in RIPA buffer (89900, Thermo Scientific) containing 1X Halt Protease and Phosphatase Inhibitor Cocktail (78440, Thermo Scientific), and equal amounts of protein (60 μg) were resolved by SDS-PAGE on a 12% acrylamide gel. Antibodies used were all obtained from Thermo Scientific and are listed in Table 4. Equal protein loading was confirmed using the Ponceau staining method.

### Animal care and husbandry

All animal experimental procedures and care were approved by the Institutional Animal Care and Use Committee. Fah^−/−^Rag2^−/−^Il2rg^−/−^ (FRG) mice were obtained from Yecuris Corporation (Tigard, Oregon), and NOD.Cg-Prkdcscid Il2rgtm1Wjl/SzJ (NSG) mice were obtained from Jackson Laboratory (Bar Harbor, ME). Six-to eight-week-old animals were kept on a 12-hour day/night cycle with free access to food and water. NSG mice were fed an irradiated mouse diet (LabDiet, 5P75) at Charles River’s Accelerator and Development Lab (Cambridge, MA). FRG mice were maintained on irradiated mouse chow with low tyrosine content (5LJ5, PicoLab®) and NTBC-containing drinking water at a concentration of 16 mg/L up until cell transplantation. NTBC was purchased from Yecuris Corporation (CuRx Nitisinone, 20-0027). The drinking water also contained prophylactic antibiotics Sulfamethoxazole at 640 µg/mL and Trimethoprim at 128 µg/mL (20-0037, Yecuris Corporation), as well as 3% dextrose water (07-869-6236, Yecuris Corporation). FRG mouse husbandry, transplantation, and collection of samples for analysis were performed at Yecuris Corporation’s testing facility (Tigard, OR).

### Transplantation of hepatocytes in FRG animals

NTBC was withdrawn from the drinking water of FRG animals the day prior to cell transplantation to initiate an NTBC on-and-off cycling scheme. An adenoviral vector expressing human urokinase (uPA, CuRx™ uPA Liver Tx Enhancer, 20-0029, Yecuris Corporation) was injected at a dose of 5E10 pfu/kg via tail vein intravenous injection 24 hours prior to transplantation. A total of 6.5E5 live PHH, EHH, EHH+, iHEP, or control cells were resuspended in 100 µL of XFAM media (Dulbecco’s Modified Eagle Medium, 10% Human Serum Albumin, 0.04 µg/mL Dexamethasone, 70 ng/mL Glucagon, and 1% Penicillin-Streptomycin). Cells were transplanted via transabdominal intrasplenic injection using standard insulin syringes with a 29-gauge needle.

Over the course of 123 days, FRG animals were subjected to several NTBC on-and-off cycles of 7-to 9-day-off/3-day-on and 11- to 18-day-off/3-day-on schedules. NTBC concentration was maintained at 8 mg/L during the on days. After 123 days, NTBC was withdrawn from the drinking water until day 163. During this extended NTBC withdrawal period, the body weights of the animals were closely monitored, and cumulative survival was measured in each group. Endpoint criteria were defined as animals reaching a body weight reduction greater than 20%, a body condition score below 2, or death.

### Transplantation of seeds in NSG animals

Seeds were either resuspended into 50 µL of media or encapsulated in 10 mm fibrin hydrogel grafts (12). Under 2% isoflurane anesthesia, seeds were transplanted into NSG mice by either intrasplenic pulp injection, kidney capsule injection, or intraperitoneal implantation of grafts onto the fat pad as previously described (13, 47, 48). In some instances, mice received intravenous low molecular weight heparin (6mg/kg, Selleck Chem, S5415) prior to surgery or intraperitoneal injection of Dexamethasone (10 mg/mL) for four consecutive days starting at the time of the transplantation and then every two days until the remainder of the study.

### Measurement of biomarkers from mouse samples

Blood was spun in SAI Infusions 300 µL capillary tubes (Serum MVCB-S-300 and K2EDTA MVCB-E-300). Hepatocyte engraftment was monitored by measuring several hepatocyte-produced proteins, including human albumin and Alpha-1-antitrypsin (A1AT), from mouse plasma using commercial ELISA kits from Bethyl Laboratories Inc. (E80-129 and E88-122) according to the manufacturer’s protocol. Levels of ammonia in plasma, as well as levels of Aspartate Aminotransferase (AST), Alanine Aminotransferase (ALT), Albumin (ALB), Total Bilirubin (TBIL), and Alkaline Phosphatase (ALP) in serum, were measured using IDEXX BioAnalytics assays.

### Infusion of seeds in Göttingen minipigs

Göttingen minipigs were procured from Marshall BioResources. The pigs were treated with an immunosuppression regimen consisting of Solu-Medrol (Methylprednisolone: 500 mg/animal IV; single administration on Day 0), Tacrolimus (0.4 mg/kg PO; daily starting on Day -5), and Orencia (abatacept-CTLA4-Ig: 15 mg/kg IV; single administration on Day 0).

On the day of seed transplantation, fasted animals were pre-anesthetized with 2– 4 mg/kg Telazol and 1–2 mg/kg Xylazine delivered by intramuscular injection. All animals received Meloxicam (0.2–0.4 mg/kg IM/PO) and Buprenorphine SR (0.1– 0.24 mg/kg SC) pre-operatively. EXCEDE was administered intramuscularly as a prophylactic antibiotic at a dose of 1.0 mL/20 kg body weight. Animals were intubated and maintained on isoflurane for effect during surgery.

All animals were placed in dorsal recumbency, and a midline incision was made through the skin, subcutaneous tissue, and linea alba. The spleen was located, gently retracted to the midline, and positioned such that the splenic artery and its arcade were identified and made visible and accessible. If needed, one of the larger branches of the splenic artery was used, where a ligature was placed proximally and distally to secure the vessel. Once secured, a catheter was placed into the artery branch. The catheter was attached to a sterile IV line primed with cells to be infused using an injection pump.

Prior to surgery, seeds were prepared by loading 1E05 seeds/kg suspended into a cryostorage bag. On the day of surgery, the cryostorage bag was placed on an IV pole, and the line was connected to the catheterized branch of the splenic artery. Seed infusion was performed at a controlled flow rate using an infusion pump (Jorgensen Lab, 1060Q). Wash media was used to resuspend the seeds in the bag during infusion and to wash the bag and IV line after the seed infusion was completed. Sodium heparin was administered intravenously during infusion at 100–300 units/kg to prevent clot formation, followed by 100–200 units/kg every 45–60 minutes.

### Histological staining of explants

Liver, graft, spleen, or kidney samples were fixed in 10% normal buffered formalin for 24 hours, washed, and stored in 70% ethanol until paraffin embedding and sectioning. Sections were stained with hematoxylin and eosin (H&E) and Picrosirius Red according to standard protocols. Engrafted hepatocytes were detected by IHC staining for FAH, human CPS1 or human CK18, and by IF staining for human OTC and human ARG1 (Table 4). All histology slides were scanned using a Leica Aperio AT2 or Ventana DP scanner.

### Quantification of human Fah staining from IHC samples

FAH staining in left lateral lobe (LLL) and median lobe (ML) liver sections was quantified using a pixel-by-pixel thresholding approach in RGB space.

Background was first eliminated by excluding pixels that exceeded a threshold for white pixels. Of the remaining pixels, in RGB color space, a clustering algorithm was used to segment pixels into either stained or unstained pixels. The stained area was calculated as stained_pixel_count / (stained_pixel_count + unstained_pixel_count).

### Human genomic DNA quantification from mouse tissue

Explanted organs were weighed, flash-frozen in liquid nitrogen, and then cryofractured using the CP02 cryoPREP pulverizer (Covaris). The pulverized tissue was homogenized overnight. DNA was extracted from the homogenate using the Maxwell RCS Tissue DNA extraction kit (Promega). Human genomic DNA was quantified using a real-time PCR method to discriminate and quantify the human DNA fraction from xenogeneic DNA, with primers and probes specific to human Alu elements (49).

Alu forward: GGTGAAACCCCGTCTCTACT

Alu reverse: GGTTCAAGCGATTCTCCTGC

Alu probe: CGCCCGGCTAATTTTTGTAT

### Bulk RNA-sequencing from cells, seeds, and mouse tissue

Gene expression from bulk RNA-sequencing was measured by incubating cells, seeds, or mouse tissue in Trizol and isolating mRNA using the PolyA capture method. The mRNA was run on a gel, and its quality was assessed with the RIN method, where a RIN > 6 was accepted. Samples were then sequenced on an Illumina HiSeq 4000 with 2x150 bp paired-end sequencing. For in vitro samples, sequences from the raw FASTQ files were aligned to GRCh38 using RSubread with default parameters (50). For in vivo samples, human and mouse transcripts were disambiguated using XenofilteR with default parameters (51). The raw counts were normalized using gene exon length and sample sequencing depth to produce transcripts per million (TPM).

### Single-cell RNA sequencing from cells

Gene expression of single cells was measured by plating all cells overnight, lifting the cells with Accutase and EDTA, and then loading them onto lanes of a 10x Genomics flowcell. The samples were sequenced with 28 bases for barcoding plus 56 bases of paired-end sequencing on an Illumina NextSeq 500 instrument running 75 cycles. Sequences in the FASTQ files were aligned to GRCh38 using RSubread with default parameters (51).

To evaluate the presence of two sub-populations, the following preprocessing pipeline was applied: (i) barcodes with fewer than 3000 reads, likely representing cell fragments, or those exceeding a proportion of mitochondrial mRNA, indicative of a dead cell based on a sample-specific cutoff from Otsu’s method (27% to 41%), were eliminated; (ii) raw counts were normalized into relative counts summing to 1 per barcode to account for differences in total reads per barcode; (iii) standard scaling (sklearn) was applied within a sample type to bring genes to a similar scale.

Following this pipeline, a Gaussian Mixture Model (sklearn) with n=2 components and the same fixed seed across all cell types was applied within a cell type. The extent of cluster separation was evaluated using the silhouette score (sklearn implementation; (52)), which measures intra-vs. inter-cluster distance. For visualization, dimensions were first reduced to n = 50 with truncated singular value decomposition (sklearn), and the output was piped to UMAP (53) with n = 2 dimensions, a fixed seed, and default parameters.

### Statistical methods

All statistical tests were performed in either Prism 10.4 or Python 3.7. Unless otherwise stated, means of two groups were compared with Student’s t-test, multiple means were compared with Tukey’s test, and survival was compared with the log-rank test. Linear regression and the significance of a univariate coefficient were evaluated with ordinary least square and a t-test comparing the standard error against the coefficient. On plots, all error bars are +/-95% confidence interval unless otherwise stated. The word “significant” was employed to represent p < 0.05.

## Acknowledgments

We thank Yecuris Corporation for their support in executing the FRG studies and Alpha Preclinical for their support in executing the minipig studies.

## Funding

This work was fully funded by Satellite Biosciences, Inc.

## Author Contributions

S.K.M., J.E.M., J.K.M., C.W., F.M., J.B., E.V., C.C., S.B., A.R., T.J.L., S.C., A.C. conceptualized and supervised the work. S.S.K., A.K.K., M.L., M.J., T.N., D.B., M.V.B, E.U., M.R., C.S., M.C. contributed to experimental design, data acquisition, and analysis. A.C. wrote the manuscript, with input and revisions from all authors. All authors reviewed and approved the final manuscript.

## Competing Interests

S.K.M., S.S.K., J.E.M., A.K.K., M.L., J.K.M., M.J., T.N., C.W., F.M., D.B., M.V.B., E.U., M.R., C.S., M.C., J.B., E.V., A.R., T.J.L., S.C., A.C. completed this work as employees of Satellite Biosciences. C.C., S.B., A.C. are founders of Satellite Biosciences. The authors have no other conflicts.

## Supplementary Information

**Figure S1:**
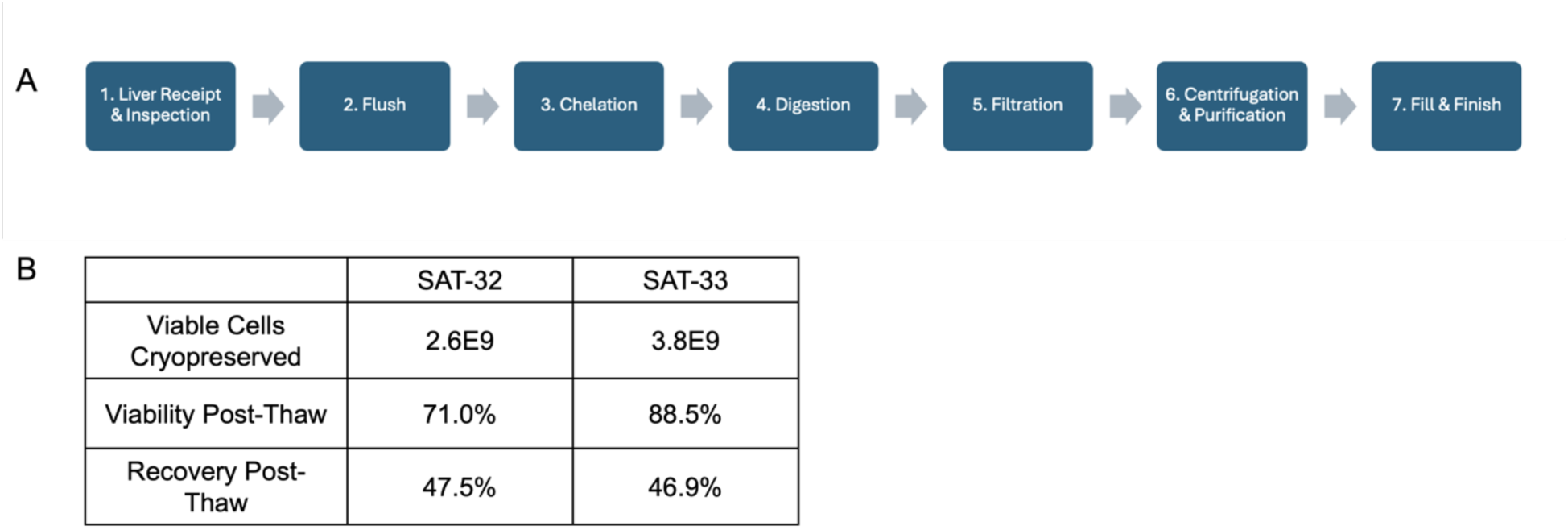
Hepatocyte isolation data. (A) Hepatocyte isolation begins with cannulating the liver’s portal vein and performing a flush to remove blood and debris. This is followed by perfusion with a chelating agent, then perfusion with collagenase-containing buffer to enzymatically digest the liver tissue, enabling hepatocyte separation. The resulting cell suspension is filtered to remove debris, and hepatocytes are purified through centrifugation. The hepatocytes are then cryopreserved. (B) Table captures data from two internally isolated lots. Of note, both donors were pediatric with livers weighing < 0.5 kg. Total viable cells cryopreserved, viability and recovery of viable cells post-thaw are listed.

**Figure S2:**
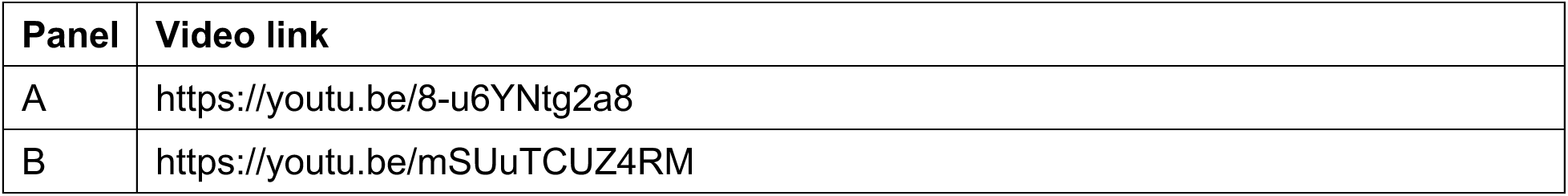
Time lapse video of P0 expansion. PHHs are plated at a low density and expanded for 13 days using expansion medium. Phase contrast pictures were taken every 24 hours, within 2 hours of same time each day. At the end of expansion, images taken on different days were stitched together to generate time lapse video. A) Overview of expansion time lapse generated using 6x6 montage for each time point. Scale bar = 2 mm. B) Zoomed in expansion time lapse generated using one image out of 36 images captured for each time point. Scale bar = 300 μm.

**Figure S3:**
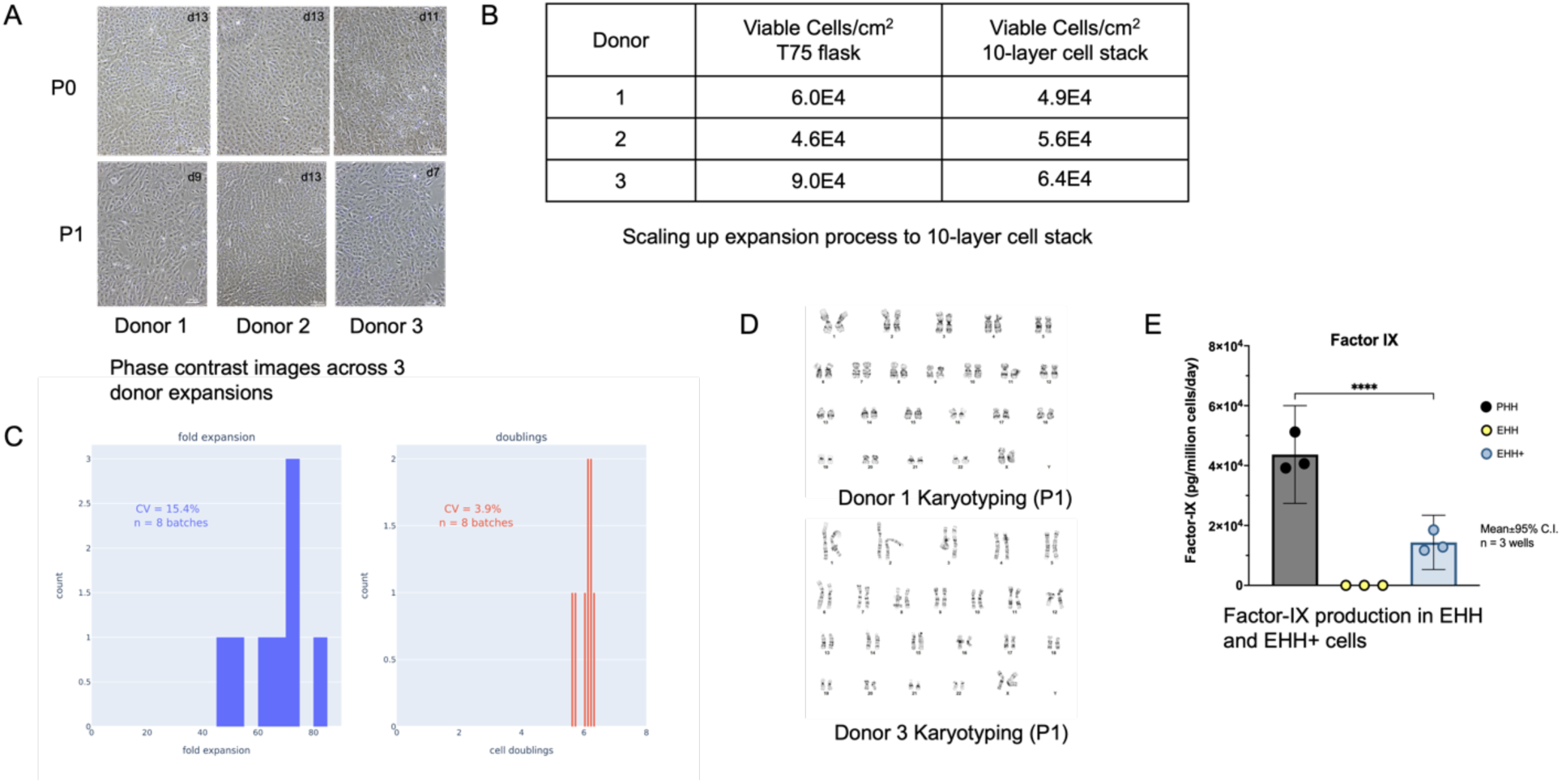
Additional characterization of EHH cells. (A) Phase contrast images of P0 and P1 expansion of Donor 1, Donor 2 and Donor 3 PHH lots using expansion protocol. Scale bar = 100 μm. (B) Scale up of expansions from T75 flasks to 10-layer CellStacks. (C) Fold expansion and cell doublings across n=8 batches generated over 1 year. Batch-to-batch CV was 15.4% for fold expansion and 3.9% for cell doublings. (D) Karyotyping analysis on P1 Donor 1 and P1 Donor 3 EHHs by GTL-banding demonstrating normal human female karyotype. No clonal aberrations were detected. (E) Levels of secreted Factor IX was quantified by ELISA in PHH, EHH and EHH+ cell supernatant and normalized to cell number at the time of supernatant collection.

**Figure S4:**
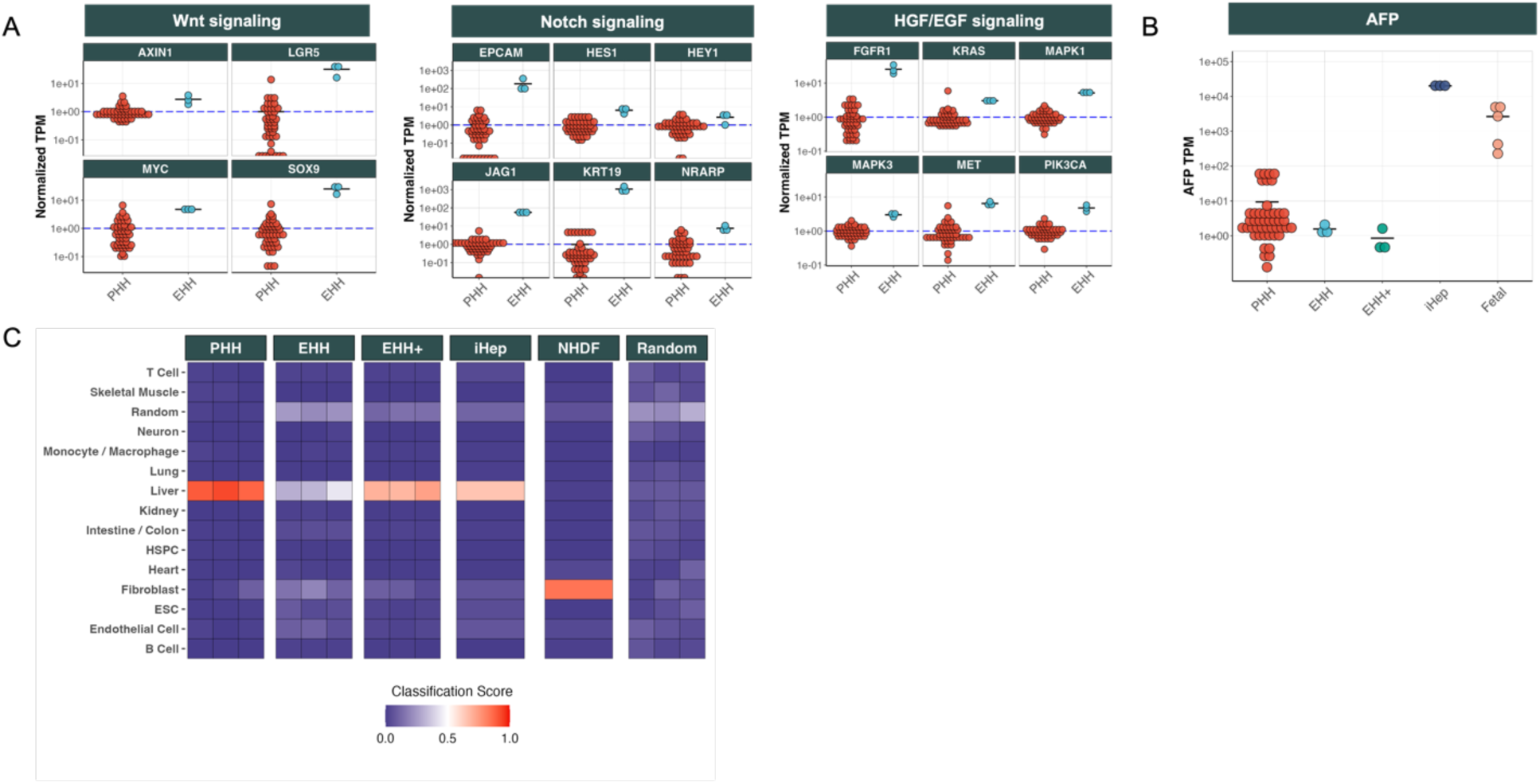
Additional RNA-seq based characterization of EHH and EHH+ cells. (A) RNA-seq based gene expression analysis demonstrating upregulation of key genes downstream of Wnt, Notch, and HGF/EGF signaling in EHHs. TPM values are normalized to the mean of the PHH group for each respective gene. (B) Alpha-fetoprotein (AFP) transcript levels (TPM) are significantly lower in PHH, EHH, and EHH+ cells compared to iHeps and fetal hepatocytes, indicating minimal reversion to a fetal-like state during expansion. (C) PACNet classification heatmap showing the classification scores of different cell types (PHH, EHH, EHH+, iHep, and NHDF). Random refers to profiles generated by random sampling of 70 cell profiles.

**Figure S5:**
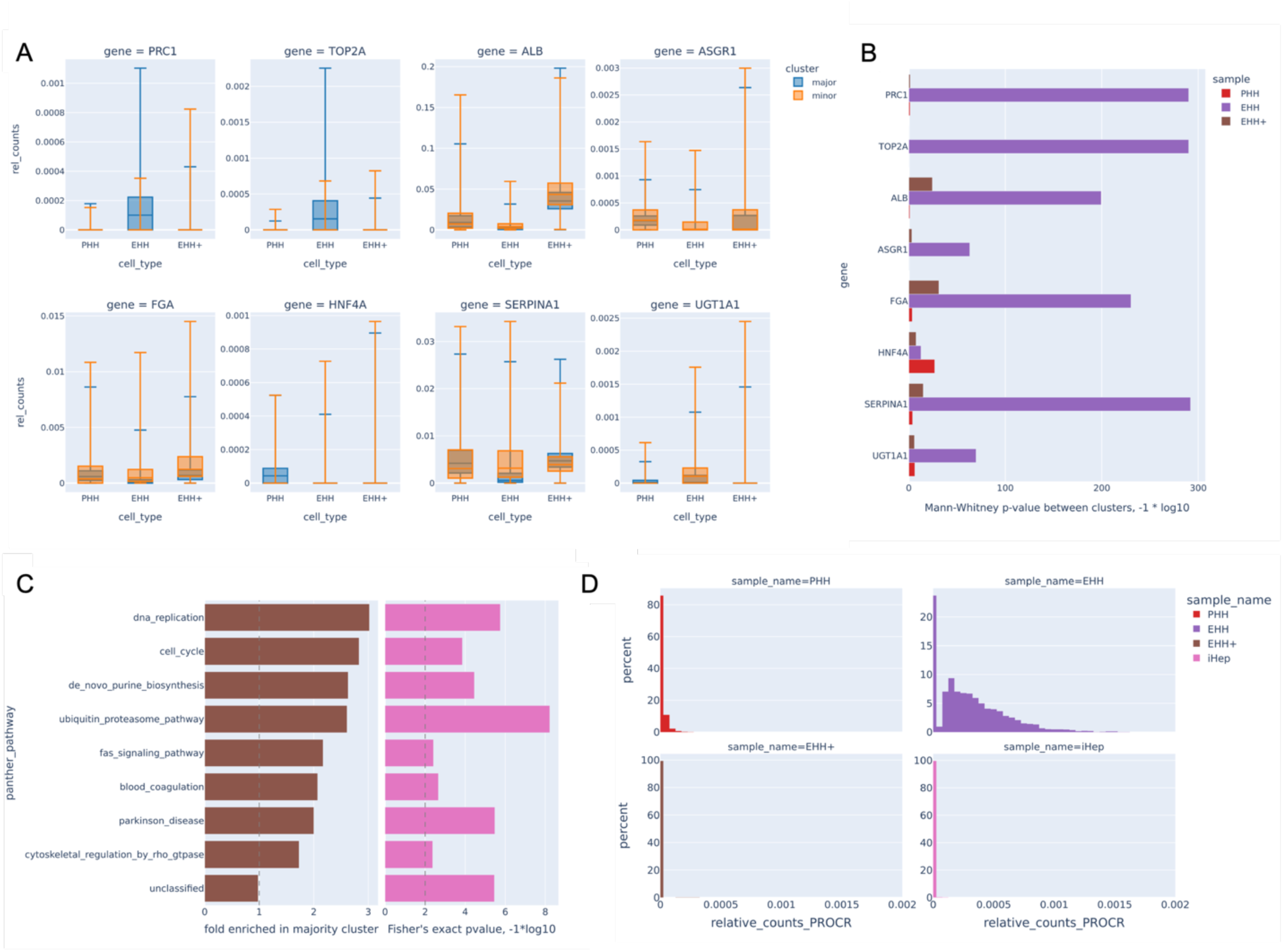
EHH cells contain a minor subpopulation that express more mature hepatic genes and less proliferative genes than the majority subpopulation. (A) scRNA-seq gene expression indicating that relative to the minority subpopulation, the majority subpopulation in EHH cells show higher expression of proliferative genes (e.g., PRC1, TOP2A) and lower expression of mature hepatic genes, which are less pronounced in PHH or EHH+ cells. (B) between the two subpopulations or clusters, based on a Mann-Whitney U-test, the largest and most consistent differences in scRNA-seq gene expression were found in EHH cells. (C) between the two EHH subpopulations, after running DESeq2 and Panther, the most enriched Panther pathways in the majority cluster were DNA replication and cell cycle, again supporting higher proliferation in the majority sub-population. (D) PROCR expression is virtually absent in PHH, EHH+, and iHep cells, but detectable in 76.3% of EHH cells.

**Figure S6:**
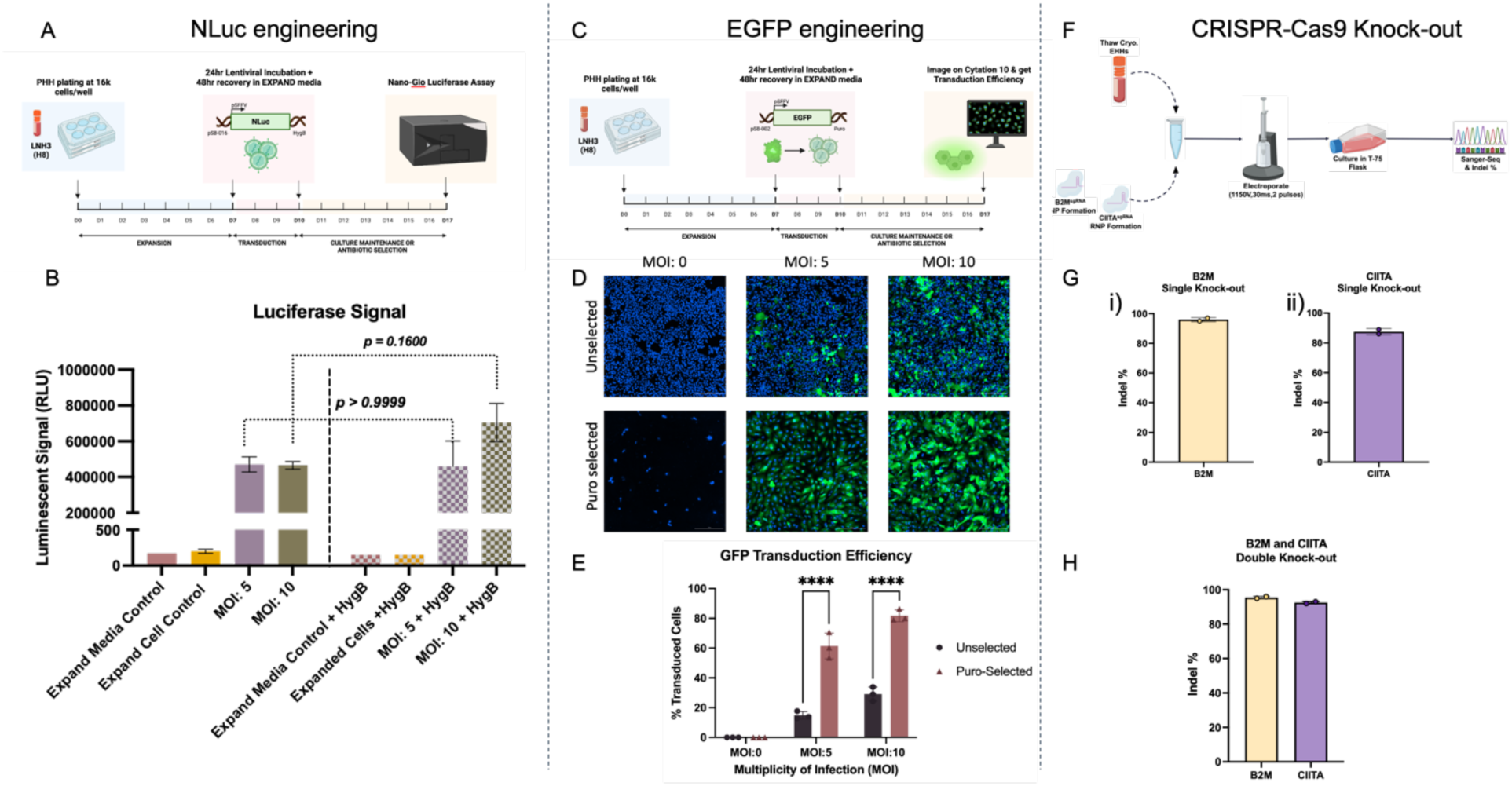
EHH cells are amenable for cell engineering involving lentivirus mediated transgene expression and CRISPR-Cas9 gene knock-out. (A) Schematic of EHH cell engineering to express NLuc transgene. PHH cells are expanded for a week before transducing with lentivirus for 24 hours followed by 48 hour recovery in Expand media. Control and transduced EHH cells are expanded between day 10 and 17 by culturing in Expand media with or without Hygromycin. (B) Luciferase activity quantified using Nano-Glo Luciferase assay demonstrating elevated signal in transduced cells (for both 5 MOI and 10 MOI, unselected and hygromycin selected samples) compared to untransduced cell control and media control. n=1 for media control conditions and n=2 for reminder of the conditions where each point represents independent well. (C) Schematic of EHH cell engineering to express EGFP transgene. PHH cells are expanded for a week before transducing with lentivirus for 24 hours followed by 48 hour recovery in Expand media. Control and transduced EHH cells are expanded between day 10 and 17 by culturing in Expand media with or without Puromycin. (D) Representative fluorescent images showing EGFP expression in EHH cells stained with Hoechst nuclear stain. Scale bar = 300 μm. (E) Percentage EGFP positive EHH cells were quantified using Cytation 10 software and plotted as a bar graph showing ∼80% EGFP positive cells in 10 MOI transduced Puromycin selected condition. n=3 where each point represents independent well. (F) Schematic showcasing the knock-out of specific targets (B2M and CIITA) in EHHs using CRISPR-Cas9 technology. Briefly, EHHs are thawed and mixed with ribonucleoproteins (RNPs) targeting B2M and/or CIITA and subsequently electroporated. The cells are then cultured in a T-75 flask with Expand media until they are harvested for downstream analytics, such as sanger sequencing and indel efficiency using inference of CRISPR edits (ICE) from Synthego. (G) Quantified indel efficiencies demonstrating high levels of genomic editing occurring in EHHs during single knock-out experiments of (i) B2M, ∼95% and (ii) CIITA, ∼90%. n=2 independent experiments per target. (H) Indel efficiencies highlighting the successful knock-out of B2M (95%) and CIITA (92%) specifically within double knock-out conditions. n=2 independent experiments.

**Figure S7:**
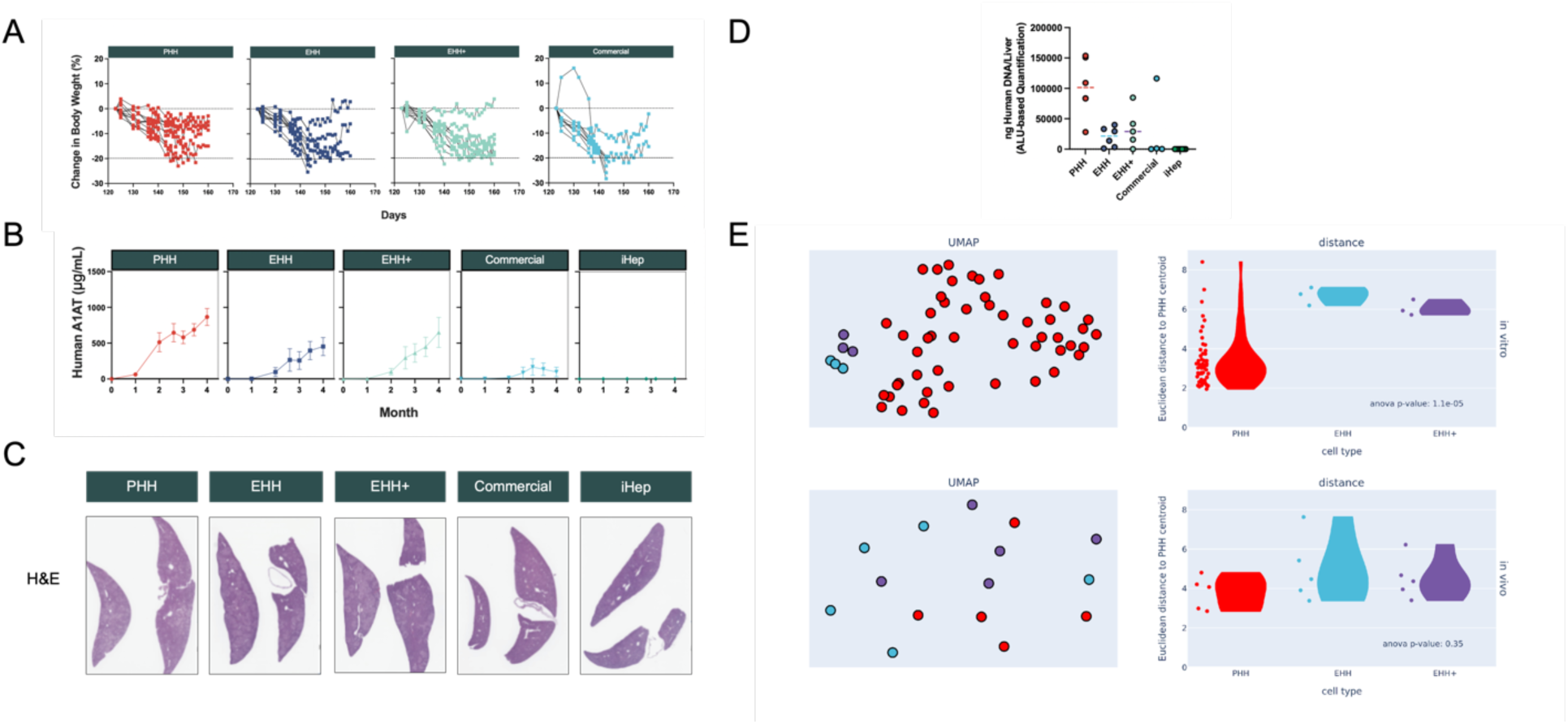
Supporting data for FRG study. FRG mice were transplanted with PHH, EHH, EHH+, Commercial or iHep cells and NTBC-cycled for 3 months (n=10-12 animals per group). After 123 days (3 months), NTBC was removed and animals followed for a period of up to 47 days. (A) Body weight change in % after NTBC removal at day 123. (B) Longitudinal Plasma Human A1AT levels. (C) Representative images of Hematoxylin and Eosin (H&E)-stained FRG mouse livers at endpoint (Magnification 1X). (D) Quantification of human DNA in mouse liver at explant by Alu-based real-time PCR method (n=5-6 per group). (E) Based on bulk RNA-seq gene expression, in vitro EHH and EHH+ cells show separation in UMAP clustering and a significantly different mean Euclidean distance to the PHH centroid relative to PHH cells (p = 1.1e-5), but this difference is abrogated once in vivo (p=0.35), presumably due to maturation of the EHH and EHH+ cells in vivo.

**Figure S8:**
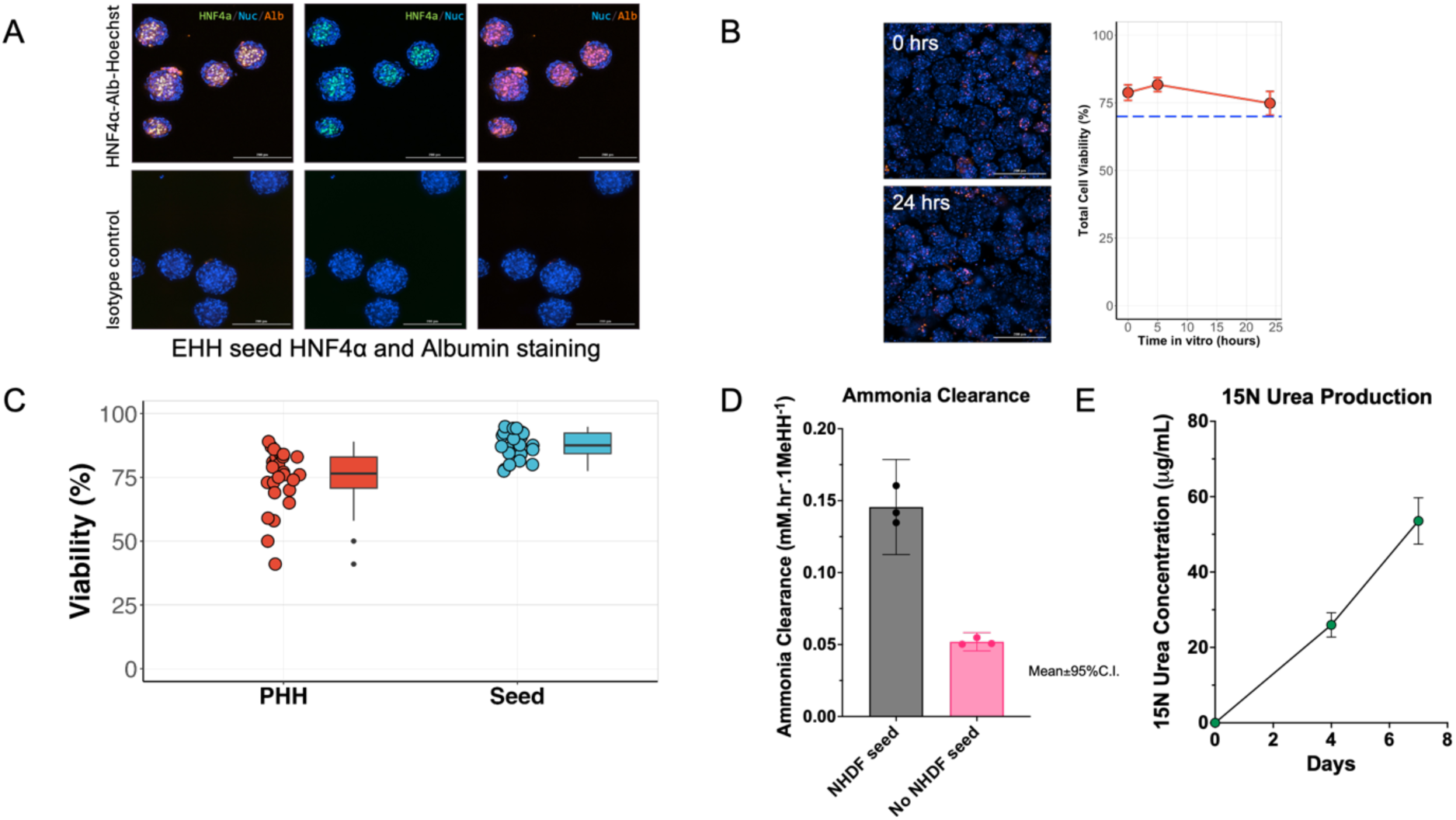
Additional EHH seed characterization. (A) Immunostaining of EHH seeds for detecting HNF4a and Albumin. Matched isotype control antibody was used to rule out non-specific signal. Nuclei were stained using Hoechst. (B) Stability of seeds over 24 hours. All cells are stained in blue (Hoechst 33342) and dead cells are stained in red (NucFix) at 0 hours and at 24 hours. Seed global cell viability remains above 70% (dashed blue line) over 24 hours in culture as assessed via an intact viability assay. Scale bars are 200 μm. (C) Comparison between EHH seed and PHH cell viability (n = 23 batches for seeds, n = 26 donors). (D) Ammonia clearance measured per well and normalized to the number of viable EHH initially seeded in each well. (E) 15N Urea concentration was determined in freshly aggregated EHH seeds and EHH seeds held in appropriate media for 4 days and 7 days. n= 2 for baseline and 7 day samples with each point representing an independent well. n=4 for 4 day samples with each point representing an independent well.

**Figure S9:**
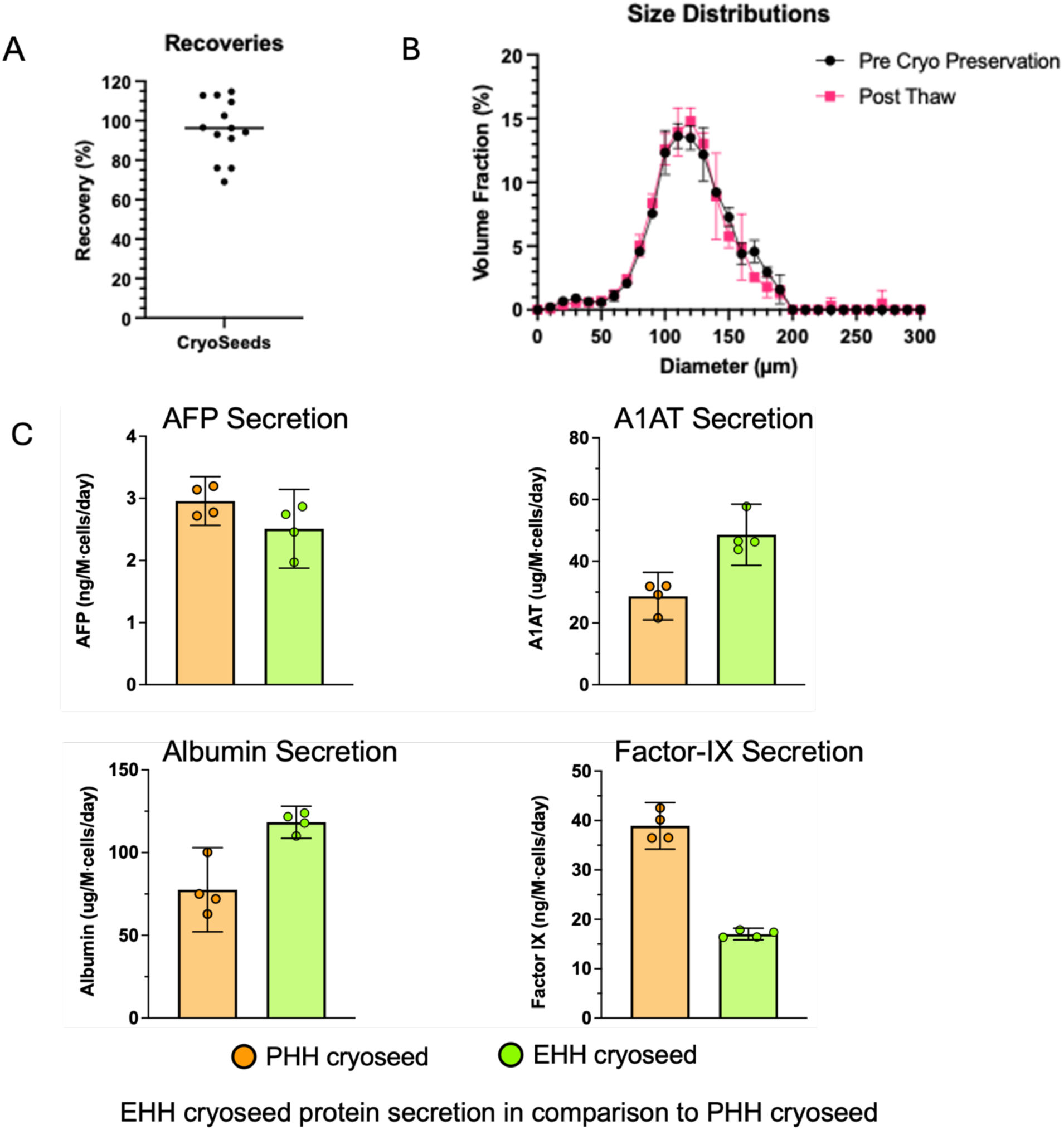
Additional characterization of cryopreserved EHH seeds. (A) Volumetric recovery percentage measured post-thaw, expressed relative to pre-cryopreservation levels. Each dot represents a unique batch. (B) Comparison of size distribution between pre-cryopreservation and post-thaw states (mean ± 95% CI, n = 4 measurements from the same batch). (C) Protein levels measured by ELISA after holding seeds for 4 days in appropriate media. Secreted protein levels normalized to the number of viable EHH cells initially seeded in each well. n=4 with each point representing an independent well.

**Figure S10:**
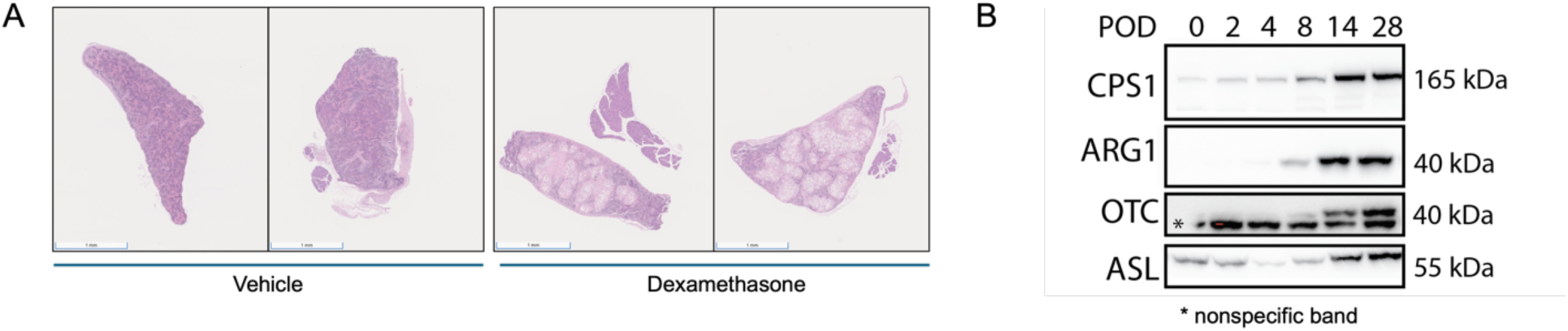
Analysis of seed engraftment with glucocorticoid administration. (A) Innate immune suppression via glucocorticoid (dexamethasone, 10 mg/kg) administration enhances engraftment within the spleen, as evidenced by an increased seed bed on H&E staining in mouse spleen samples at POD28. (B) Western blot analysis of liver-specific proteins in the spleen reveals progressive functional maturation of EHH seeds in vivo over 28 days with glucocorticoid administration (10 mg/kg).

**Figure S11:**
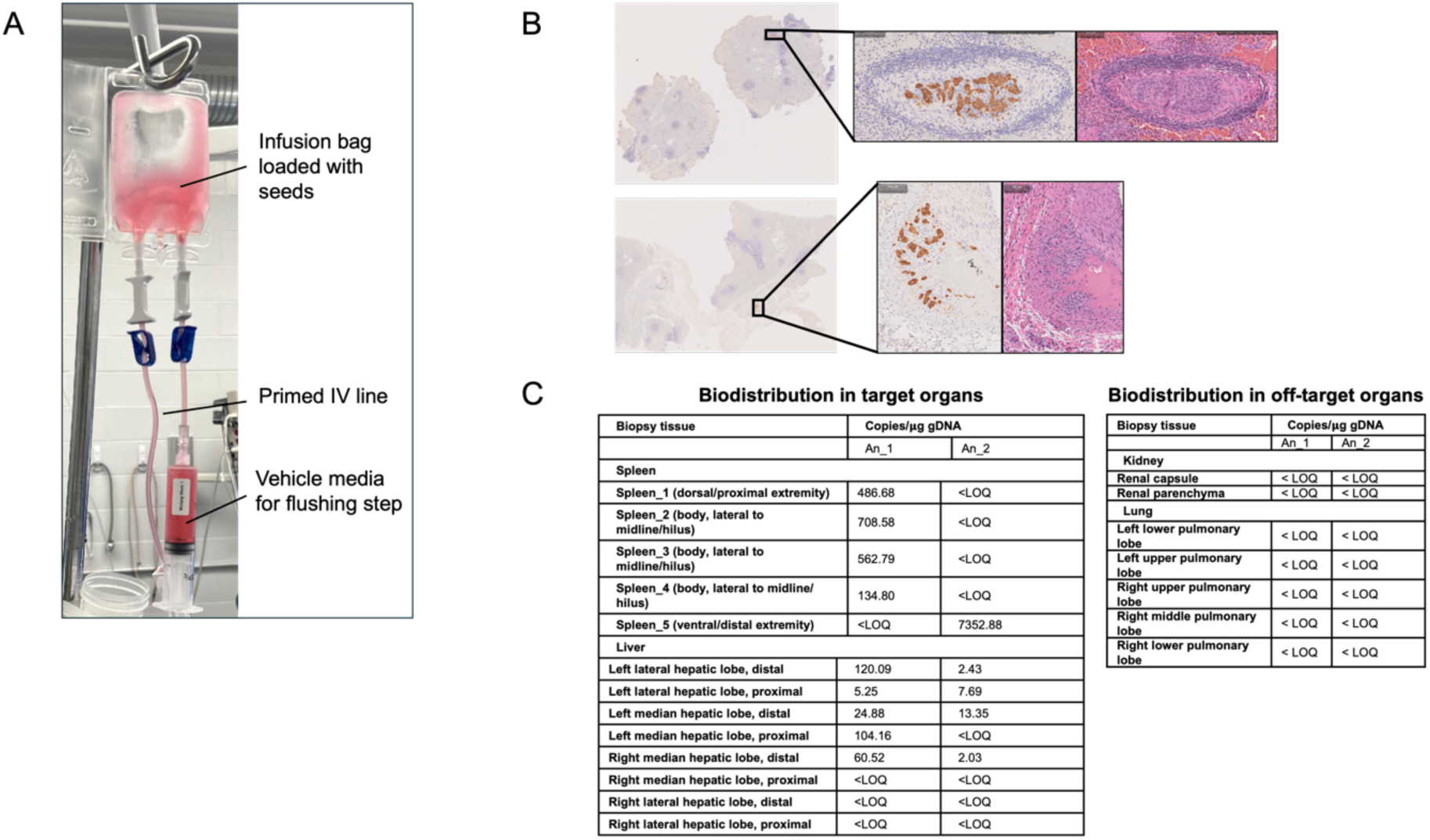
Minipig seed delivery. (A) Cryostorage bag setup for seed infusion. (B) Presence of seeds is confirmed by CK18 (left) and hematoxylin/eosin (right) staining of spleen tissue samples at POD2. (C) The biodistribution of seeds in target organs reveals predominant localization in the spleen, with some distribution to the liver (LOQ = 1 copy/ µg gDNA).

## References

1. FastStats - Chronic Liver Disease or Cirrhosis. Available at: https://www.cdc.gov/nchs/fastats/liver-disease.htm?utm_source=chatgpt.com [Accessed 25 December 2024].

2. B. J. Dwyer, M. T. Macmillan, P. N. Brennan, S. J. Forbes, Cell therapy for advanced liver diseases: Repair or rebuild. J. Hepatol. 74, 185–199 (2021).

3. J. Nulty, H. Anand, A. Dhawan, Human Hepatocyte Transplantation: Three Decades of Clinical Experience and Future Perspective. Stem Cells Transl. Med. 13, 204–218 (2023).

4. A. Shlomai, et al., Modeling host interactions with hepatitis B virus using primary and induced pluripotent stem cell-derived hepatocellular systems. Proc. Natl. Acad. Sci. 111, 12193–12198 (2014).

5. E. Michailidis, et al., Expansion, in vivo–ex vivo cycling, and genetic manipulation of primary human hepatocytes. Proc. Natl. Acad. Sci. 117, 1678– 1688 (2020).

6. I. J. Fox, J. Roy-Chowdhury, Hepatocyte transplantation. J. Hepatol. 40, 878– 886 (2004).

7. V. Sauer, N. Roy-Chowdhury, C. Guha, J. Roy-Chowdhury, Induced Pluripotent Stem Cells as a Source of Hepatocytes. Curr. Pathobiol. Rep. 2, 11– 20 (2014).

8. X. Gao, et al., Hepatocyte-like cells derived from human induced pluripotent stem cells using small molecules: implications of a transcriptomic study. Stem Cell Res. Ther. 11, 393 (2020).

9. P. Huang, et al., Direct Reprogramming of Human Fibroblasts to Functional and Expandable Hepatocytes. Cell Stem Cell 14, 370–384 (2014).

10. C. Wang, et al., Dedifferentiation-associated inflammatory factors of long-term expanded human hepatocytes exacerbate their elimination by macrophages during liver engraftment. Hepatology 76, 1690–1705 (2022).

11. S. Ogawa, et al., Three-dimensional culture and cAMP signaling promote the maturation of human pluripotent stem cell-derived hepatocytes. Development 140, 3285–3296 (2013).

12. T. Takebe, et al., Vascularized and functional human liver from an iPSC-derived organ bud transplant. Nature 499, 481–484 (2013).

13. K. R. Stevens, et al., In situ expansion of engineered human liver tissue in a mouse model of chronic liver disease. Sci Transl Med 9, eaah5505 (2017).

14. F. Wu, et al., Generation of hepatobiliary organoids from human induced pluripotent stem cells. J. Hepatol. 70, 1145–1158 (2019).

15. H. Ma, et al., The nuclear receptor THRB facilitates differentiation of human PSCs into more mature hepatocytes. Cell Stem Cell 29, 795–809.e11 (2022).

16. E. Fitzpatrick, et al., Coculture with Mesenchymal Stem Cells Results in Improved Viability and Function of Human Hepatocytes. Cell Transplant. 24, 73– 83 (2013).

17. K. A. Soltys, et al., Barriers to the successful treatment of liver disease by hepatocyte transplantation. J. Hepatol. 53, 769–774 (2010).

18. A. A. Chen, et al., Humanized mice with ectopic artificial liver tissues. Proc National Acad Sci 108, 11842–11847 (2011).

19. A. X. Chen, et al., Controlled Apoptosis of Stromal Cells to Engineer Human Microlivers. Adv Funct Mater 30, 1910442 (2020).

20. A. Chhabra, et al., A vascularized model of the human liver mimics regenerative responses. Proc National Acad Sci 119 (2022).

21. E. K. W. Lo, et al., Platform-agnostic CellNet enables cross-study analysis of cell fate engineering protocols. Stem Cell Rep. 18, 1721–1742 (2023).

22. T. Rathbone, et al., Electroporation-Mediated Delivery of Cas9 Ribonucleoproteins Results in High Levels of Gene Editing in Primary Hepatocytes. CRISPR J. 5, 397–409 (2022).

23. H. Azuma, et al., Robust expansion of human hepatocytes in Fah−/−/Rag2−/−/Il2rg−/− mice. Nat. Biotechnol. 25, 903–910 (2007).

24. K. Zhang, et al., In Vitro Expansion of Primary Human Hepatocytes with Efficient Liver Repopulation Capacity. Cell Stem Cell 23, 806–819.e4 (2018).

25. S. Zhu, et al., Mouse liver repopulation with hepatocytes generated from human fibroblasts. Nature 508, 93–97 (2014).

26. Y. Gao, et al., Distinct Gene Expression and Epigenetic Signatures in Hepatocyte-like Cells Produced by Different Strategies from the Same Donor. Stem Cell Rep. 9, 1813–1824 (2017).

27. L. T. Ang, et al., A Roadmap for Human Liver Differentiation from Pluripotent Stem Cells. Cell Rep. 22, 2190–2205 (2018).

28. M. J. Caldez, M. Bjorklund, P. Kaldis, Cell cycle regulation in NAFLD: when imbalanced metabolism limits cell division. Hepatol. Int. 14, 463–474 (2020).

29. H. Hu, et al., Long-Term Expansion of Functional Mouse and Human Hepatocytes as 3D Organoids. Cell 175, 1591–1606.e19 (2018).

30. D. Wang, et al., Long-Term Expansion of Pancreatic Islet Organoids from Resident Procr+ Progenitors. Cell 180, 1198–1211.e19 (2020).

31. J. L. Corbett, S. A. Duncan, iPSC-Derived Hepatocytes as a Platform for Disease Modeling and Drug Discovery. Front. Med. 6, 265 (2019).

32. J. Blaszkiewicz, S. A. Duncan, Advancements in Disease Modeling and Drug Discovery Using iPSC-Derived Hepatocyte-like Cells. Genes 13, 573 (2022).

33. I. Tamargo-Rubio, A. B. Simpson, J. A. Hoogerland, J. Fu, Human induced pluripotent stem cell–derived liver-on-a-chip for studying drug metabolism: the challenge of the cytochrome P450 family. Front. Pharmacol. 14, 1223108 (2023).

34. A. Laemmle, et al., Aquaporin 9 induction in human iPSC-derived hepatocytes facilitates modeling of ornithine transcarbamylase deficiency. Hepatology 76, 646–659 (2022).

35. C. Xiang, et al., Long-term functional maintenance of primary human hepatocytes in vitro. Science 364, 399–402 (2019).

36. S. R. Khetani, S. N. Bhatia, Microscale culture of human liver cells for drug development. Nat Biotechnol 26, 120–126 (2008).

37. F. N. Smets, Y. Chen, L.-J. Wang, H. E. Soriano, Loss of cell anchorage triggers apoptosis (anoikis) in primary mouse hepatocytes. Mol. Genet. Metab. 75, 344–352 (2002).

38. K. Zhang, et al., Efficient expansion and CRISPR-Cas9-mediated gene correction of patient-derived hepatocytes for treatment of inherited liver diseases. Cell Stem Cell 31, 1187–1202.e8 (2024).

39. C. J. Flaim, S. Chien, S. N. Bhatia, An extracellular matrix microarray for probing cellular differentiation. Nat Methods 2, 119–125 (2005).

40. J. C. Y. Dunn, M. L. Yarmush, H. G. Koebe, R. G. Tompkins, Hepatocyte function and extracellular matrix geometry: long-term culture in a sandwich configuration. FASEB J. 3, 174–177 (1989).

41. S. N. Bhatia, U. J. Balis, M. L. Yarmush, M. Toner, Effect of cell–cell interactions in preservation of cellular phenotype: cocultivation of hepatocytes and nonparenchymal cells. Faseb J 13, 1883–1900 (1999).

42. S. R. Khetani, A. A. Chen, B. Ranscht, S. N. Bhatia, T-cadherin modulates hepatocyte functions in vitro. FASEB J. 22, 3768–3775 (2008).

43. E. E. Hui, S. N. Bhatia, Micromechanical control of cell–cell interactions. Proc. National Acad. Sci. 104, 5722–5726 (2007).

44. X. Liu, et al., Isolation of primary human liver cells from normal and nonalcoholic steatohepatitis livers. STAR Protoc. 4, 102391 (2023).

45. Y. Chen, et al., A versatile polypharmacology platform promotes cytoprotection and viability of human pluripotent and differentiated cells. Nat. Methods 18, 528–541 (2021).

46. N. Otsu, A Threshold Selection Method from Gray-Level Histograms. *IEEE Trans. Syst., Man*, Cybern. 9, 62–66 (1979).

47. K. P. O’Rourke, et al., Transplantation of engineered organoids enables rapid generation of metastatic mouse models of colorectal cancer. Nat. Biotechnol. 35, 577–582 (2017).

48. N. Matsumoto, et al., Techniques of fragile renal organoids transplantation in mice. Acta Cirúrgica Bras. 36, e361102 (2021).

49. S. Funakoshi, et al., Enhanced engraftment, proliferation and therapeutic potential in heart using optimized human iPSC-derived cardiomyocytes. Sci. Rep. 6, 19111 (2016).

50. Y. Liao, G. K. Smyth, W. Shi, The R package Rsubread is easier, faster, cheaper and better for alignment and quantification of RNA sequencing reads. Nucleic Acids Res 47, gkz114-(2019).

51. R. J. C. Kluin, et al., XenofilteR: computational deconvolution of mouse and human reads in tumor xenograft sequence data. Bmc Bioinformatics 19, 366 (2018).

52. P. J. Rousseeuw, Silhouettes: A graphical aid to the interpretation and validation of cluster analysis. J. Comput. Appl. Math. 20, 53–65 (1987).

53. L. McInnes, J. Healy, J. Melville, UMAP: Uniform Manifold Approximation and Projection for Dimension Reduction. Arxiv (2018).

