## Supplementary Information for "Expandable, Functional Hepatocytes Derived from Primary Cells Enable Liver Therapeutics"

A

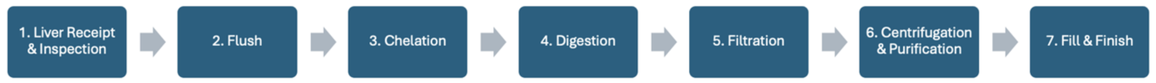


B

| | SAT-32 | SAT-33 |
|---|---|---|
| Viable Cells Cryopreserved | 2.6E9 | 3.8E9 |
| Viability Post-Thaw | 71.0% | 88.5% |
| Recovery Post-Thaw | 47.5% | 46.9% |

**Figure S1: Hepatocyte isolation data.** (A) Hepatocyte isolation begins with cannulating the liver's portal vein and performing a flush to remove blood and debris. This is followed by perfusion with a chelating agent, then perfusion with collagenase-containing buffer to enzymatically digest the liver tissue, enabling hepatocyte separation. The resulting cell suspension is filtered to remove debris, and hepatocytes are purified through centrifugation. The hepatocytes are then cryopreserved. (B) Table captures data from two internally isolated lots. Of note, both donors were pediatric with livers weighing < 0.5 kg. Total viable cells cryopreserved, viability and recovery of viable cells post-thaw are listed.

| Panel | Video link |
| --- | --- |
| A | https://youtu.be/8-u6YNtg2a8 |
| B | https://youtu.be/mSUuTCUZ4RM |

**Figure S2: Time lapse video of P0 expansion.** PHHs are plated at a low density and expanded for 13 days using expansion medium. Phase contrast pictures were taken every 24 hours, within 2 hours of same time each day. At the end of expansion, images taken on different days were stitched together to generate time lapse video. A) Overview of expansion time lapse generated using 6x6 montage for each time point. Scale bar = 2 mm. B) Zoomed in expansion time lapse generated using one image out of 36 images captured for each time point. Scale bar = 300 μm.

A

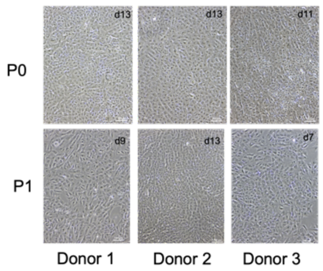


Phase contrast images across 3 donor expansions

B

| Donor | Viable Cells/cm$^2$ T75 flask | Viable Cells/cm$^2$ 10-layer cell stack |
|---|---|---|
| 1 | 6.0E4 | 4.9E4 |
| 2 | 4.6E4 | 5.6E4 |
| 3 | 9.0E4 | 6.4E4 |

Scaling up expansion process to 10-layer cell stack

C

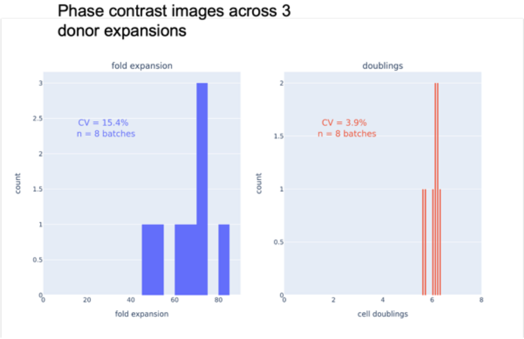


D

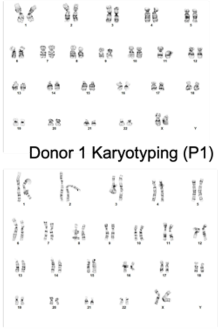

Donor 1 Karyotyping (P1)

Donor 3 Karyotyping (P1)

E

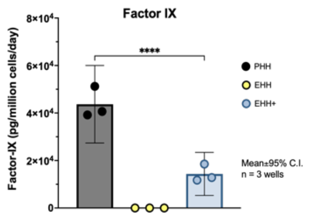


Factor-IX production in EHH and EHH+ cells

**Figure S3: Additional characterization of EHH cells.** (A) Phase contrast images of P0 and P1 expansion of Donor 1, Donor 2 and Donor 3 PHH lots using expansion protocol. Scale bar = 100 µm. (B) Scale up of expansions from T75 flasks to 10-layer CellStacks. (C) Fold expansion and cell doublings across n=8 batches generated over 1 year. Batch-to-batch CV was 15.4% for fold expansion and 3.9% for cell doublings. (D) Karyotyping analysis on P1 Donor 1 and P1 Donor 3 EHHs by GTL-banding demonstrating normal human female karyotype. No clonal aberrations were detected. (E) Levels of secreted Factor IX was quantified by ELISA in PHH, EHH and EHH+ cell supernatant and normalized to cell number at the time of supernatant collection.

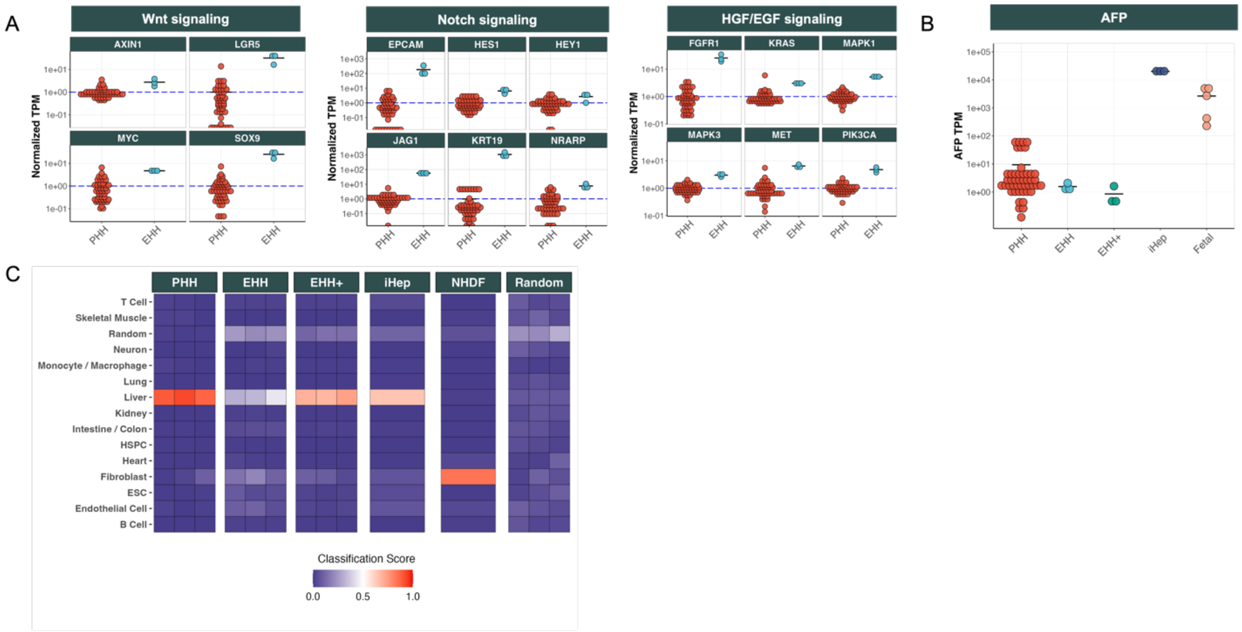


**Figure S4: Additional RNA-seq based characterization of EHH and EHH+ cells.** (A) RNA-seq based gene expression analysis demonstrating upregulation of key genes downstream of Wnt, Notch, and HGF/EGF signaling in EHHs. TPM values are normalized to the mean of the PHH group for each respective gene. (B) Alpha-fetoprotein (AFP) transcript levels (TPM) are significantly lower in PHH, EHH, and EHH+ cells compared to iHeps and fetal hepatocytes, indicating minimal reversion to a fetal-like state during expansion. (C) PACNet classification heatmap showing the classification scores of different cell types (PHH, EHH, EHH+, iHep, and NHDF). Random refers to profiles generated by random sampling of 70 cell profiles.

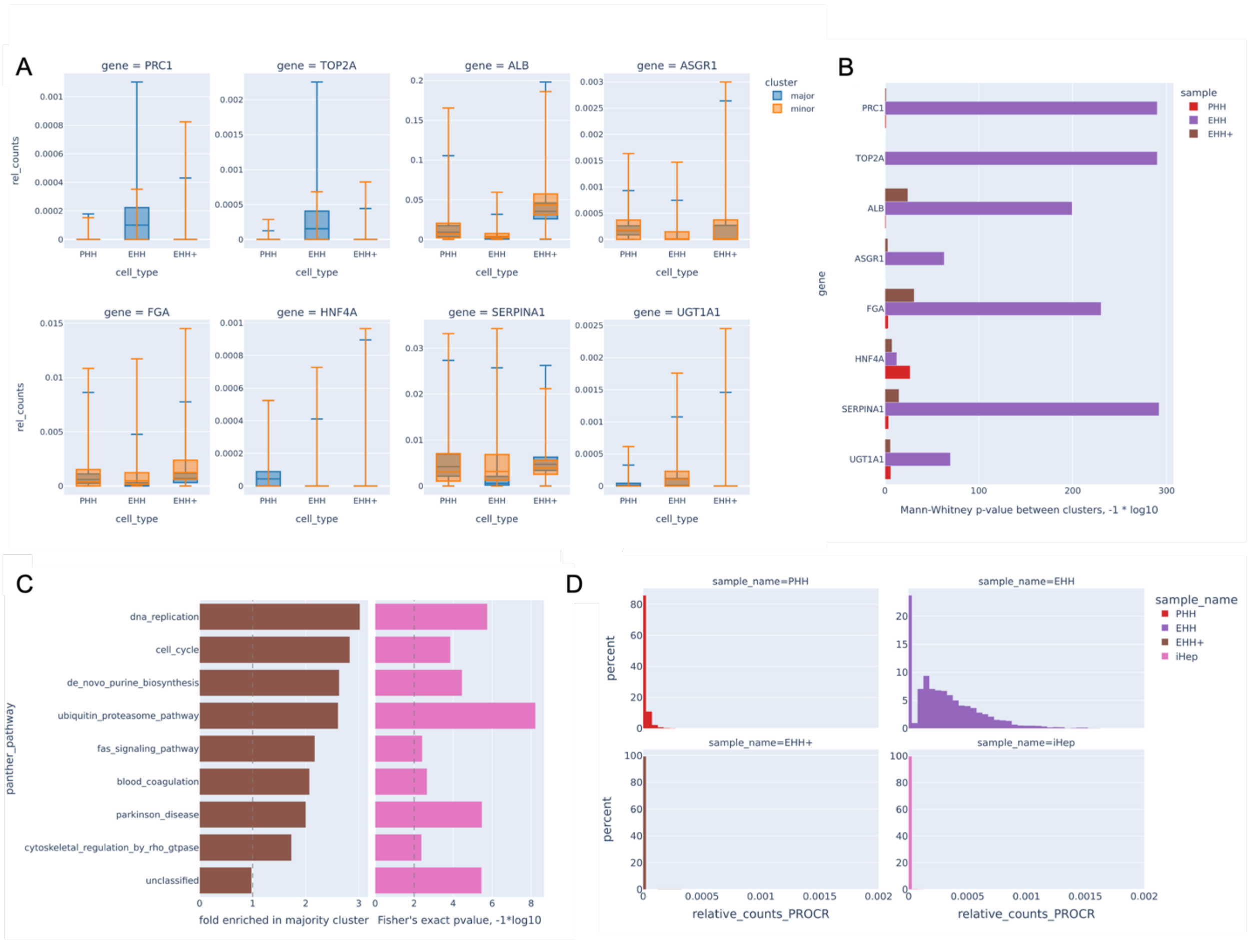

**Figure S5: EHH cells contain a minor subpopulation that express more mature hepatic genes and less proliferative genes than the majority subpopulation.** (A) scRNA-seq gene expression indicating that relative to the minority subpopulation, the majority subpopulation in EHH cells show higher expression of proliferative genes (e.g., PRC1, TOP2A) and lower expression of mature hepatic genes, which are less pronounced in PHH or EHH+ cells. (B) between the two subpopulations or clusters, based on a Mann-Whitney U-test, the largest and most consistent differences in scRNA-seq gene expression were found in EHH cells. (C) between the two EHH subpopulations, after running DESeq2 and Panther, the most enriched Panther pathways in the majority cluster were DNA replication and cell cycle, again supporting higher proliferation in the majority sub-population. (D) PROCR expression is virtually absent in PHH, EHH+, and iHep cells, but detectable in 76.3% of EHH cells.

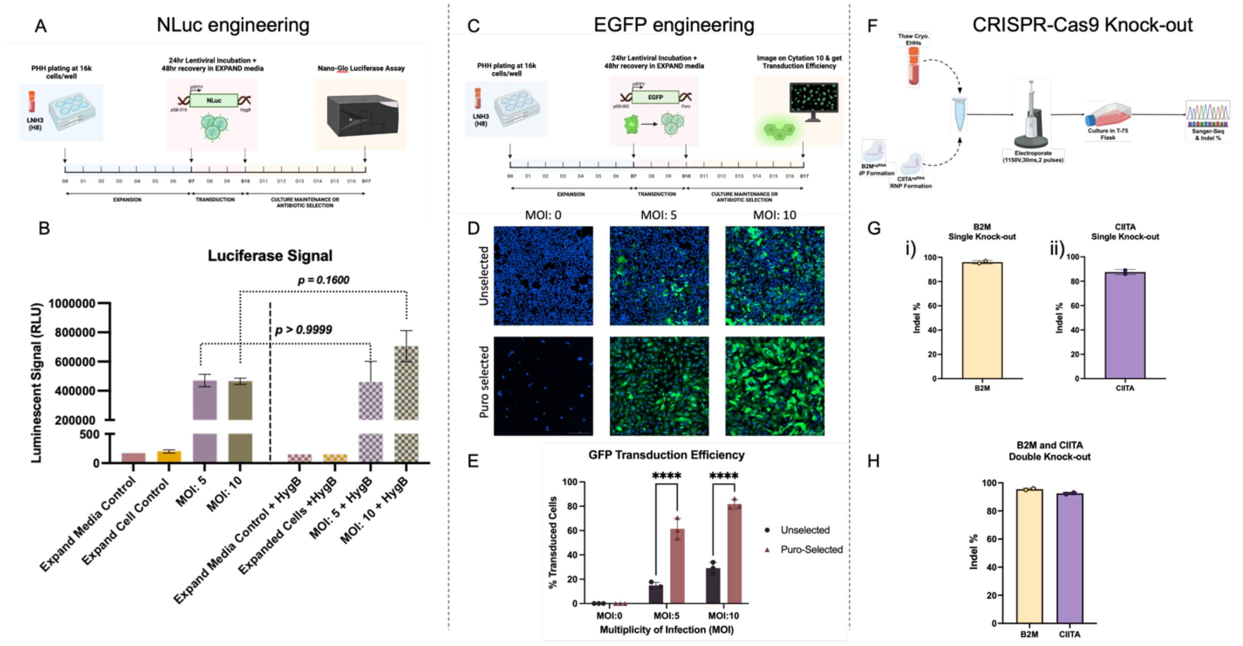


**Figure S6: EHH cells are amenable for cell engineering involving lentivirus mediated transgene expression and CRISPR-Cas9 gene knock-out.** (A) Schematic of EHH cell engineering to express NLuc transgene. PHH cells are expanded for a week before transducing with lentivirus for 24 hours followed by 48 hour recovery in Expand media. Control and transduced EHH cells are expanded between day 10 and 17 by culturing in Expand media with or without Hygromycin. (B) Luciferase activity quantified using Nano-Glo Luciferase assay demonstrating elevated signal in transduced cells (for both 5 MOI and 10 MOI, unselected and hygromycin selected samples) compared to untransduced cell control and media control. n=1 for media control conditions and n=2 for reminder of the conditions where each point represents independent well. (C) Schematic of EHH cell engineering to express EGFP transgene. PHH cells are expanded for a week before transducing with lentivirus for 24 hours followed by 48 hour recovery in Expand media. Control and transduced EHH cells are expanded between day 10 and 17 by culturing in Expand media with or without Puromycin. (D) Representative fluorescent images showing EGFP expression in EHH cells stained with Hoechst nuclear stain. Scale bar = 300 µm. (E) Percentage EGFP positive EHH cells were quantified using Cytation 10 software and plotted as a bar graph showing ~80% EGFP positive cells in 10 MOI transduced Puromycin selected condition. n=3 where each point represents independent well. (F) Schematic showcasing the knock-out of specific targets (B2M and CIITA) in EHHs using CRISPR-Cas9 technology. Briefly, EHHs are thawed and mixed with ribonucleoproteins (RNPs) targeting B2M and/or CIITA and subsequently electroporated. The cells are then cultured in a T-75 flask with Expand media until they are harvested for downstream analytics, such as sanger sequencing and indel efficiency using inference of CRISPR edits (ICE) from Synthego. (G) Quantified indel efficiencies demonstrating high levels of genomic editing occurring in EHHs during single knock-out experiments of (i) B2M, ~95% and (ii) CIITA, ~90%. n=2 independent experiments per target. (H) Indel efficiencies highlighting the successful knock-out of B2M (95%) and CIITA (92%) specifically within double knock-out conditions. n=2 independent experiments.

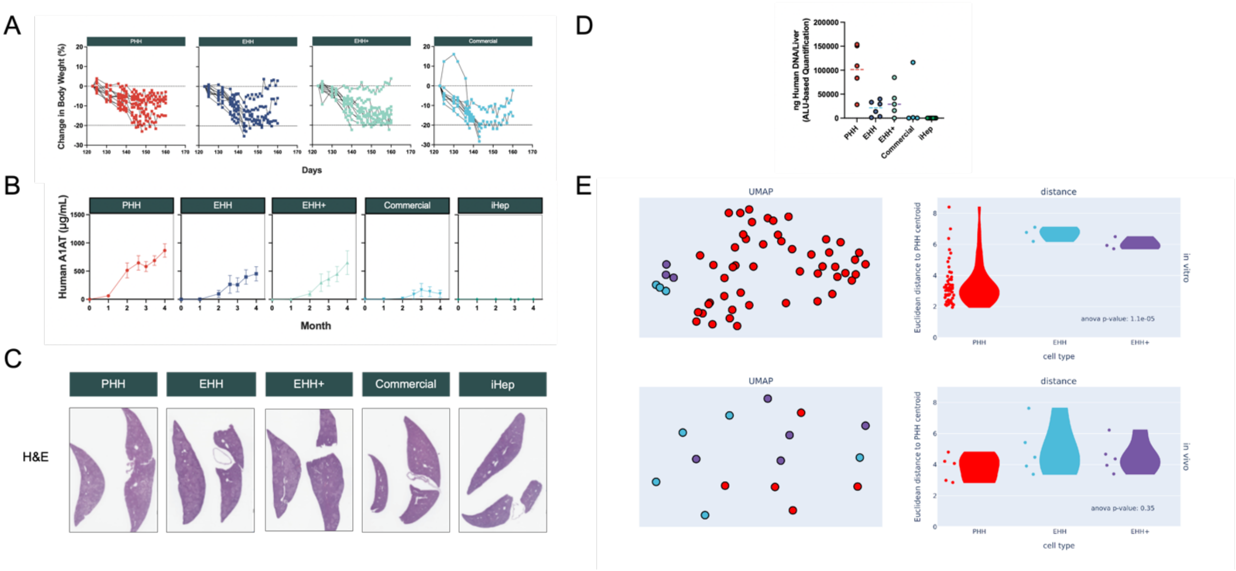


**Figure S7: Supporting data for FRG study.** FRG mice were transplanted with PHH, EHH, EHH+, Commercial or iHep cells and NTBC-cycled for 3 months (n=10-12 animals per group). After 123 days (3 months), NTBC was removed and animals followed for a period of up to 47 days. (A) Body weight change in % after NTBC removal at day 123. (B) Longitudinal Plasma Human A1AT levels. (C) Representative images of Hematoxylin and Eosin (H&E)-stained FRG mouse livers at endpoint (Magnification 1X). (D) Quantification of human DNA in mouse liver at explant by Alu-based real-time PCR method (n=5-6 per group). (E) Based on bulk RNA-seq gene expression, in vitro EHH and EHH+ cells show separation in UMAP clustering and a significantly different mean Euclidean distance to the PHH centroid relative to PHH cells (p = 1.1e-5), but this difference is abrogated once in vivo (p=0.35), presumably due to maturation of the EHH and EHH+ cells in vivo.

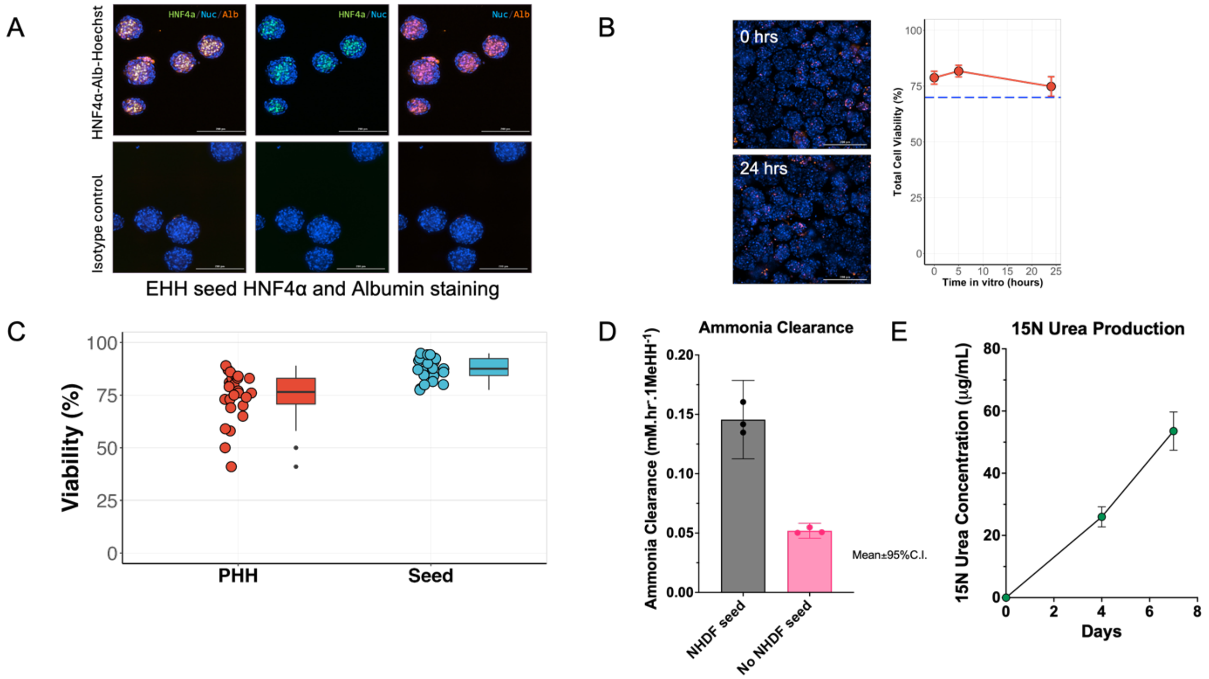


**Figure S8: Additional EHH seed characterization.** (A) Immunostaining of EHH seeds for detecting HNF4a and Albumin. Matched isotype control antibody was used to rule out non-specific signal. Nuclei were stained using Hoechst. (B) Stability of seeds over 24 hours. All cells are stained in blue (Hoechst 33342) and dead cells are stained in red (NucFix) at 0 hours and at 24 hours. Seed global cell viability remains above 70% (dashed blue line) over 24 hours in culture as assessed via an intact viability assay. Scale bars are 200 μm. (C) Comparison between EHH seed and PHH cell viability (n = 23 batches for seeds, n = 26 donors). (D) Ammonia clearance measured per well and normalized to the number of viable EHH initially seeded in each well. (E) 15N Urea concentration was determined in freshly aggregated EHH seeds and EHH seeds held in appropriate media for 4 days and 7 days. n= 2 for baseline and 7 day samples with each point representing an independent well. n=4 for 4 day samples with each point representing an independent well.

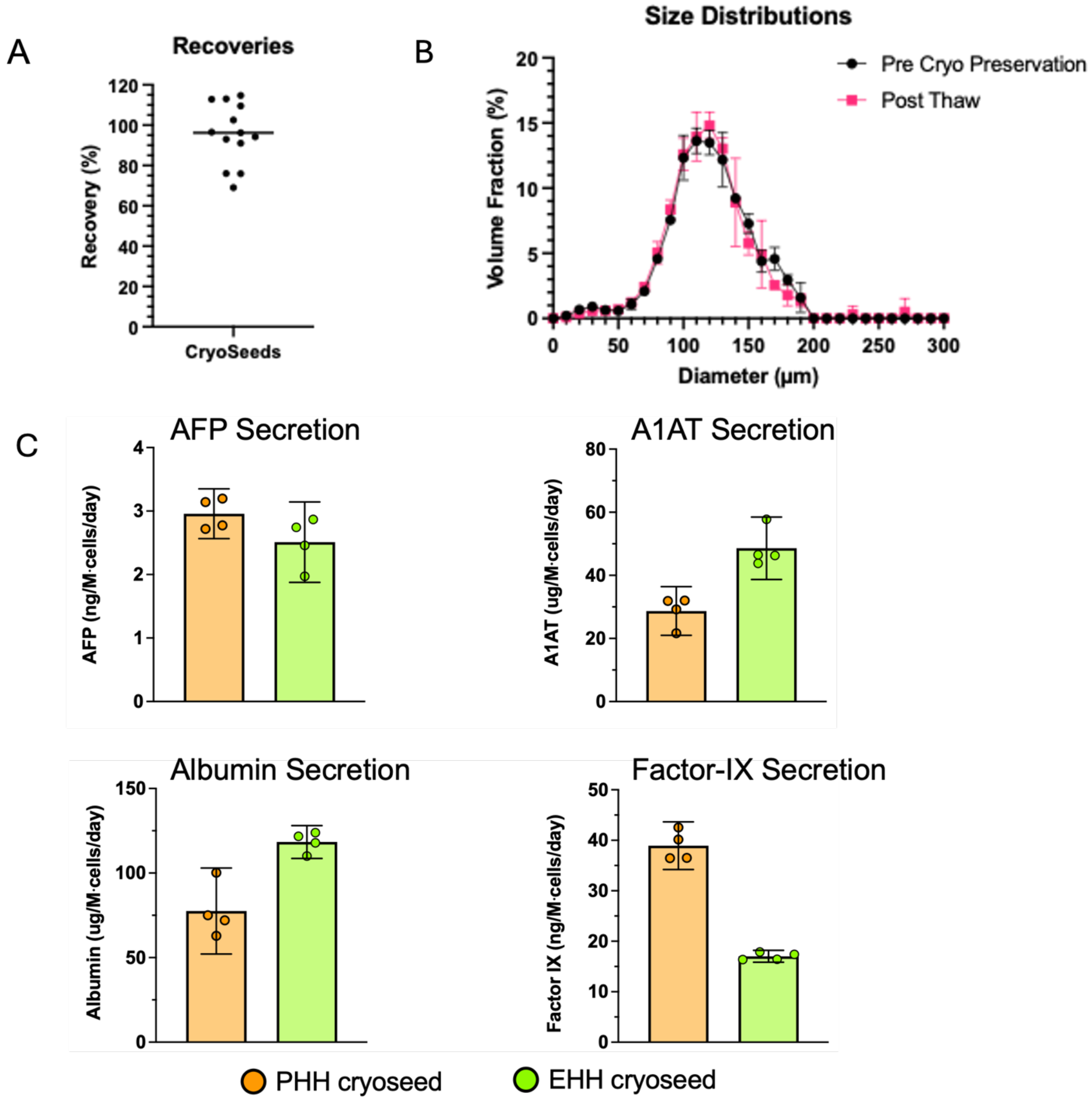


EHH cryoseed protein secretion in comparison to PHH cryoseed

**Figure S9: Additional characterization of cryopreserved EHH seeds.** (A) Volumetric recovery percentage measured post-thaw, expressed relative to pre-cryopreservation levels. Each dot represents a unique batch. (B) Comparison of size distribution between pre-cryopreservation and post-thaw states (mean ± 95% CI, n = 4 measurements from the same batch). (C) Protein levels measured by ELISA after holding seeds for 4 days in appropriate media. Secreted protein levels normalized to the number of viable EHH cells initially seeded in each well. n=4 with each point representing an independent well.

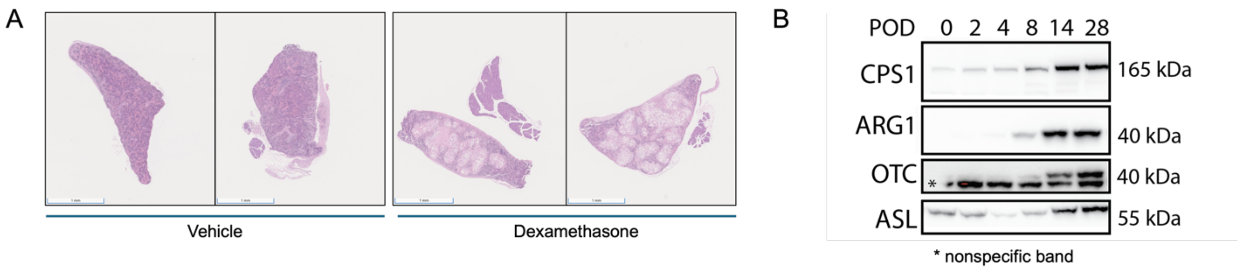


**Figure S10: Analysis of seed engraftment with glucocorticoid administration.** (A) Innate immune suppression via glucocorticoid (dexamethasone, 10 mg/kg) administration enhances engraftment within the spleen, as evidenced by an increased seed bed on H&E staining in mouse spleen samples at POD28. (B) Western blot analysis of liver-specific proteins in the spleen reveals progressive functional maturation of EHH seeds in vivo over 28 days with glucocorticoid administration (10 mg/kg).

A

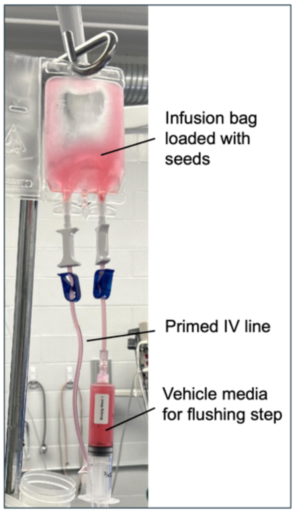


B

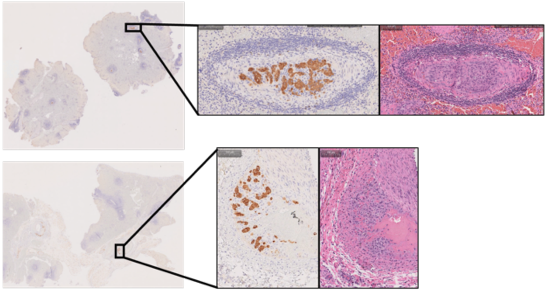

C

**Biodistribution in target organs**

| Biopsy tissue | Copies/µg gDNA | |
|---|---|---|
| | An_1 | An_2 |
| Spleen | | |
| Spleen_1 (dorsal/proximal extremity) | 486.68 | <LOQ |
| Spleen_2 (body, lateral to midline/hilus) | 708.58 | <LOQ |
| Spleen_3 (body, lateral to midline/hilus) | 562.79 | <LOQ |
| Spleen_4 (body, lateral to midline/ hilus) | 134.80 | <LOQ |
| Spleen_5 (ventral/distal extremity) | <LOQ | 7352.88 |
| Liver | | |
| Left lateral hepatic lobe, distal | 120.09 | 2.43 |
| Left lateral hepatic lobe, proximal | 5.25 | 7.69 |
| Left median hepatic lobe, distal | 24.88 | 13.35 |
| Left median hepatic lobe, proximal | 104.16 | <LOQ |
| Right median hepatic lobe, distal | 60.52 | 2.03 |
| Right median hepatic lobe, proximal | <LOQ | <LOQ |
| Right lateral hepatic lobe, distal | <LOQ | <LOQ |
| Right lateral hepatic lobe, proximal | <LOQ | <LOQ |

**Biodistribution in off-target organs**

| Biopsy tissue | Copies/µg gDNA | |
|---|---|---|
| | An_1 | An_2 |
| Kidney | | |
| Renal capsule | < LOQ | < LOQ |
| Renal parenchyma | < LOQ | < LOQ |
| Lung | | |
| Left lower pulmonary lobe | < LOQ | < LOQ |
| Left upper pulmonary lobe | < LOQ | < LOQ |
| Right upper pulmonary lobe | < LOQ | < LOQ |
| Right middle pulmonary lobe | < LOQ | < LOQ |
| Right lower pulmonary lobe | < LOQ | < LOQ |

**Figure S11: Minipig seed delivery.** (A) Cryostorage bag setup for seed infusion. (B) Presence of seeds is confirmed by CK18 (left) and hematoxylin/eosin (right) staining of spleen tissue samples at POD2. (C) The biodistribution of seeds in target organs reveals predominant localization in the spleen, with some distribution to the liver (LOQ = 1 copy/ µg gDNA).
